# A biophysical regulator of inhibitory integration and learning in mesolimbic dopamine neurons

**DOI:** 10.1101/344499

**Authors:** Kauê M. Costa, Niklas Hammer, Christopher Knowlton, Jochen Schwenk, Tamara Müller, Dorothea Schulte, Bernd Fakler, Carmen C. Canavier, Jochen Roeper

**Author notes:** National Institute on Drug Abuse Intramural Research Program, National Institutes of Health, Baltimore, USA.

## Abstract

Midbrain dopamine neurons are essential for flexible control of adaptive behaviors. DA neurons that project to different target regions have unique biophysical properties, and it is thought that this diversity reflects functional specialization. This assumption implies the presence of specific genetic determinants with precise impacts on behavior. We tested this general hypothesis by homing in on one particular biophysical mechanism, Kv4 channel inactivation, using a combination of molecular, proteomic, electrophysiological, computational, and behavioral approaches. We demonstrate that KChIP4a, a singular Kv4 β-subunit splice variant, prolongs hyperpolarization-rebound delays selectively in dopamine neurons projecting to the nucleus accumbens core, shifts the integration of inhibitory inputs and, in turn, selectively regulates learning from negative prediction-errors. Our results reveal a highly specialized, gene-to-behavior mechanistic chain that is only operative in a particular dopaminergic subsystem, illuminating how molecularly defined biophysical switches are employed for neuron subtype-specific information processing in the brain.

**Graphical abstract:** 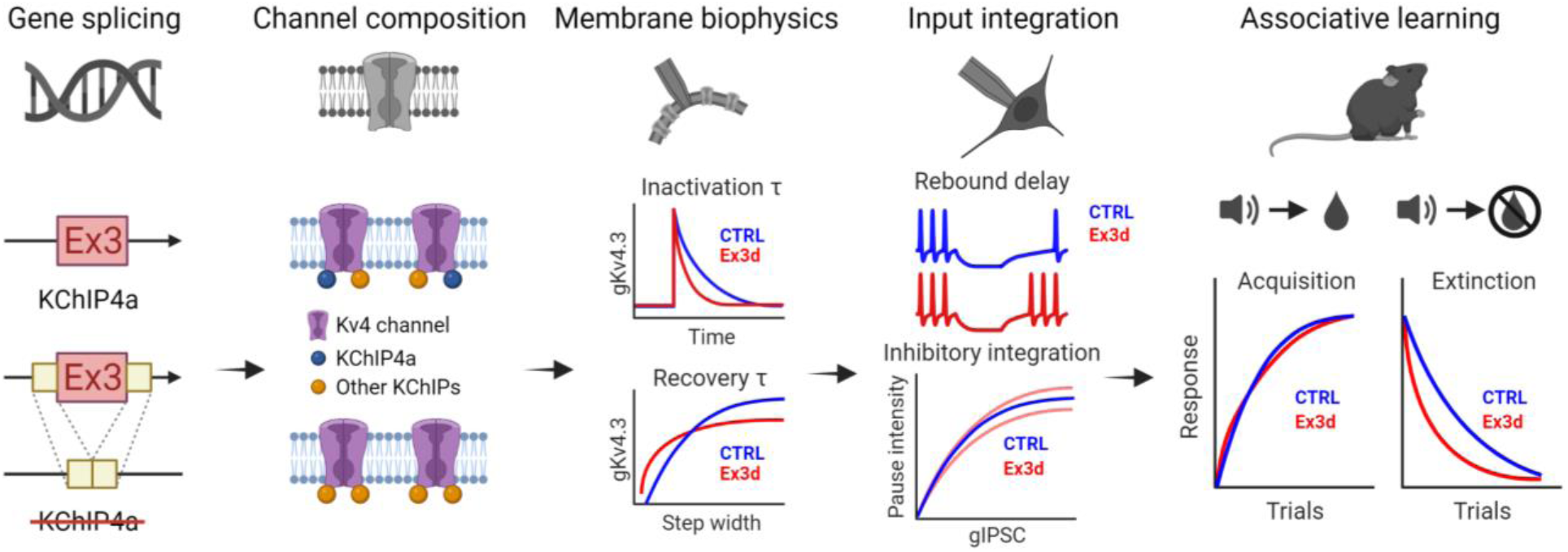

## Introduction

Midbrain dopamine (DA) neurons regulate several behavioral functions, particularly learning and motivation, and are implicated in nearly every mental illness ^1,2^. While previously thought to be a homogenous population, work from our lab and others has revealed that these neurons display a wide range of biophysical properties and gene expression profiles, which vary categorically depending on their axonal projection target and define vulnerability to disease. One hypothesis for this variability is that the membrane properties of each subgroup are specialized to a particular behavioral function ^3^. If this is true, it stands to reason that there should also be specific genetic mechanisms within each subgroup that underpin these adaptations. This would also predict that some biophysical variables, as they contribute to identifiable computational processes, should have relatively circumscribed influences in behavior. In this study, we test this general hypothesis by homing in on one key biophysical differentiator of DA neuron subpopulations.

Specifically, DA neurons in the ventral tegmental area (VTA) that project to the core of the nucleus accumbens (DA-cNAcc), among select others, display longer rebound delays from hyperpolarizing inhibition than those that project to the lateral shell of the NAcc (DA-lNacc) and the dorsolateral striatum (DA-DLS) ^4^. Atypical, slowly inactivating A-type K^+^ currents, mediated by Kv4.3 channels, are responsible for these longer delays ^5,6^. This suggests that Kv4 channels in DA-cNAcc neurons may be differentially modulated by a so far unknown mechanism.

Kv4 channel function is tightly controlled by modulatory beta subunits, including those of the KChIP (K^+^ channel interacting proteins) family ^7,8^. In mammals, four genes (KCNIP 1-4) encode distinct KChIP proteins, each with multiple splice variants. One of those, KChIP4a (a KChIP4 isoform) stands out for having a unique K^+^ channel inhibitory domain (KID), which markedly slows Kv4 channel inactivation while promoting close-state inactivation ^7,9,10^. Most KChIP isoforms, including all other KChIP4 variants, induce the opposite effects, and the ultimate impact on channel kinetics depends on the relative ratio between KChIP4a and other KChIP isoforms within the channel complex ^11,12^. Differential expression of KChIP4a by DA-cNAcc neurons is expected to produce longer post-inhibitory rebound delays by slowing inactivation of Kv4 channels. This keeps the cells not only in a more hyperpolarized voltage range for longer periods, but also alters their synaptic integration by increasing their input conductances. We therefore reasoned that KChIP4a could be one of the molecular determinants of the projection-specific diversity in rebound delays across DA neurons.

What would this mean for behavior? It has been demonstrated that the activity of many DA neurons tracks reward prediction errors (RPEs), i.e. the difference between expected and experienced value ^1^. Specifically, these DA neurons respond with phasic increases in firing to unexpected rewards and reward-predicting cues (positive RPEs), and with phasic decreases in firing, or pauses, when an expected reward is omitted (negative RPEs) ^1,13^. RPE encoding is particularly prevalent in DA neurons that project to the cNAcc ^14,15^, and multiple studies have also demonstrated that transient inhibition of DA neuron firing is causally implicated in learning from negative RPEs ^16–18^. Therefore, we predicted that the endogenous expression of the alternative splice variant KChIP4a, by controlling the integration of inhibitory inputs in DA-cNAcc neurons, might selectively regulate learning from negative RPEs.

We tested this set of hypotheses by creating a cell type- and exon-specific transgenic mouse line where the KChIP4a isoform is deleted only in DA neurons, allowing us to disrupt this proposed mechanistic chain from channel subunit to behavior at its most basic link. Using a combination of molecular, proteomic, electrophysiological, computational, and behavioral approaches, we show that KChIP4a underlies one of the key features of DA neuron diversity, and that it regulates the integration of inhibitory inputs and learning from negative RPEs.

## Results

### Selective deletion of KChIP4a from midbrain DA neurons

To selectively knock out KChIP4a in DA neurons, we developed a mouse line where we floxed the exon coding for the KID of KCNIP4 (exon 3), which is unique to the KChIP4a variant (Figure 1A and B) ^9^. We crossed these mice with a DAT-Cre line to create mice with a selective deletion of KChip4a in midbrain DA neurons (Ex3d, for “exon 3 deletion”, Figure 1C). For all subsequent procedures, these mice were compared with DAT-Cre^+/-^ littermate controls (CTRL), and both male and female mice were used for all experiments ^19^.

**Figure 1.**
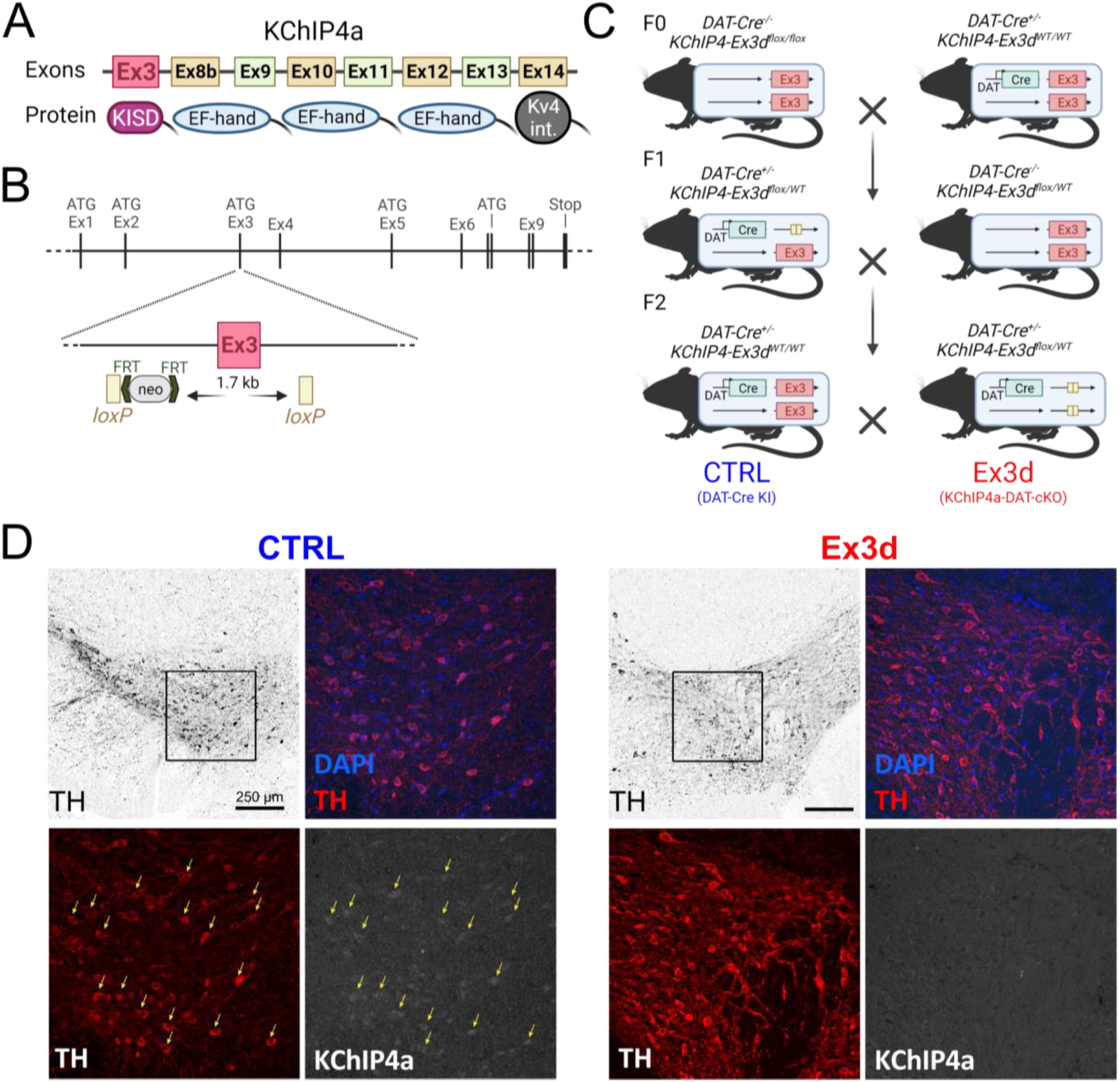
Transgenic strategy for selectively deleting KChIP4a from DA neurons. **A:** Cartoon schematic of KChIP4a exons and corresponding proteins. Note that the KChIP4a-defining KID is encoded by Ex3 of the KCNIP4 gene. **B:** Cartoon schematic of the transgenic strategy used for creating the vector where Ex3 was floxed. This vector was later electroporated into mouse embryonic stem cells which were subsequently used to generate mice with a Cre-dependent conditional KO of KCNIP4 Ex3 (see Methods section for details). **C:** Breeding strategy for generating mice with a selective deletion of KChip4a in midbrain DA neurons (Ex3d). Conditional KO mice were crossed with DAT-Cre KI mice (F0), generating double heterozygous offspring (F1), which were then crossed to produce Ex3d mice and controls that were DAT-Cre KI but did not have floxed KCNIP Ex3 (CTRL). **D:** FISH validation of the KO strategy. CTRL mice showed robust expression of KChIP4a mRNA in midbrain DA neurons, while Ex3d mice showed no such expression (see Figure S1 for additional validation). N=3 for Ex3d and CTRL.

To validate the effects of our conditional exon-specific knockout, we combined fluorescent *in situ* hybridization (FISH) with probes targeting either the KID n-terminal sequence (specific to KChIP4a) or the conserved KChIP4 c-terminus (to label all KChIP4 variants), with immunohistochemistry (IHC) for tyrosine hydroxylase (TH) to identify DA neurons. We observed clear double-labeling for both KChIp4a mRNA and TH in the midbrain of CTRL mice, but no such labeling was seen in Ex3d mice (Figure 1D). Labeling of KChIP4 mRNA was visible in TH-cells in both CTRL and Ex3d mice, confirming that our mutation ablated KChIP4a mRNA expression only in TH+ DA neurons (Figure S1J). Finally, when using the general KChIP4 probe, we observed labeling in TH+ cells in both CTRL and Ex3d mice (Figures S1E and K), demonstrating that the Ex3d mutation did not eliminate expression of other KChIP4 variants in DA neurons.

### Ex3d mutation changes midbrain Kv4 channel composition

To assess the effects of the Ex3d mutation on Kv4 channel composition, we performed quantitative high-resolution proteomic analysis of the Kv4 channel complexes (Figures 2 and S2) affinity-isolated from tissue samples that were excised from the ventral midbrain (region of interest) or cerebellum (control) of both CTRL and Ex3d mice ^20–22^. We found that the average Kv4 channel core in the ventral midbrain is an asymmetric tetramer composed of one Kv4.2 and three Kv4.3 subunits. In the midbrain, this channel core co-assembled with an average four KChIP subunits with relative contribution of 0.5 KChIP1, 0.5 KChIP2, one KChIP3 and two KChIP4 thus uncovering/revealing KChIP4 as the dominant auxiliary subunit in that region (Figure 2A). In the cerebellum, the relative contribution of KChIP4 was decreased in favor of KChIPs 1 and 3, thus identifying the high relative abundance of KChIP4 in the midbrain as region-specific (Figure 2B). Importantly, there was no significant difference in the relative abundance of all Kv4 complex constituents between CTRL and Ex3d mice, in both midbrain and cerebellum. This confirmed that the Ex3d mutation does neither disrupt expression of other KChIP4 variants, nor does it affect the overall stoichiometry of the native Kv4-KchIP channel complexes.

**Figure 2.**
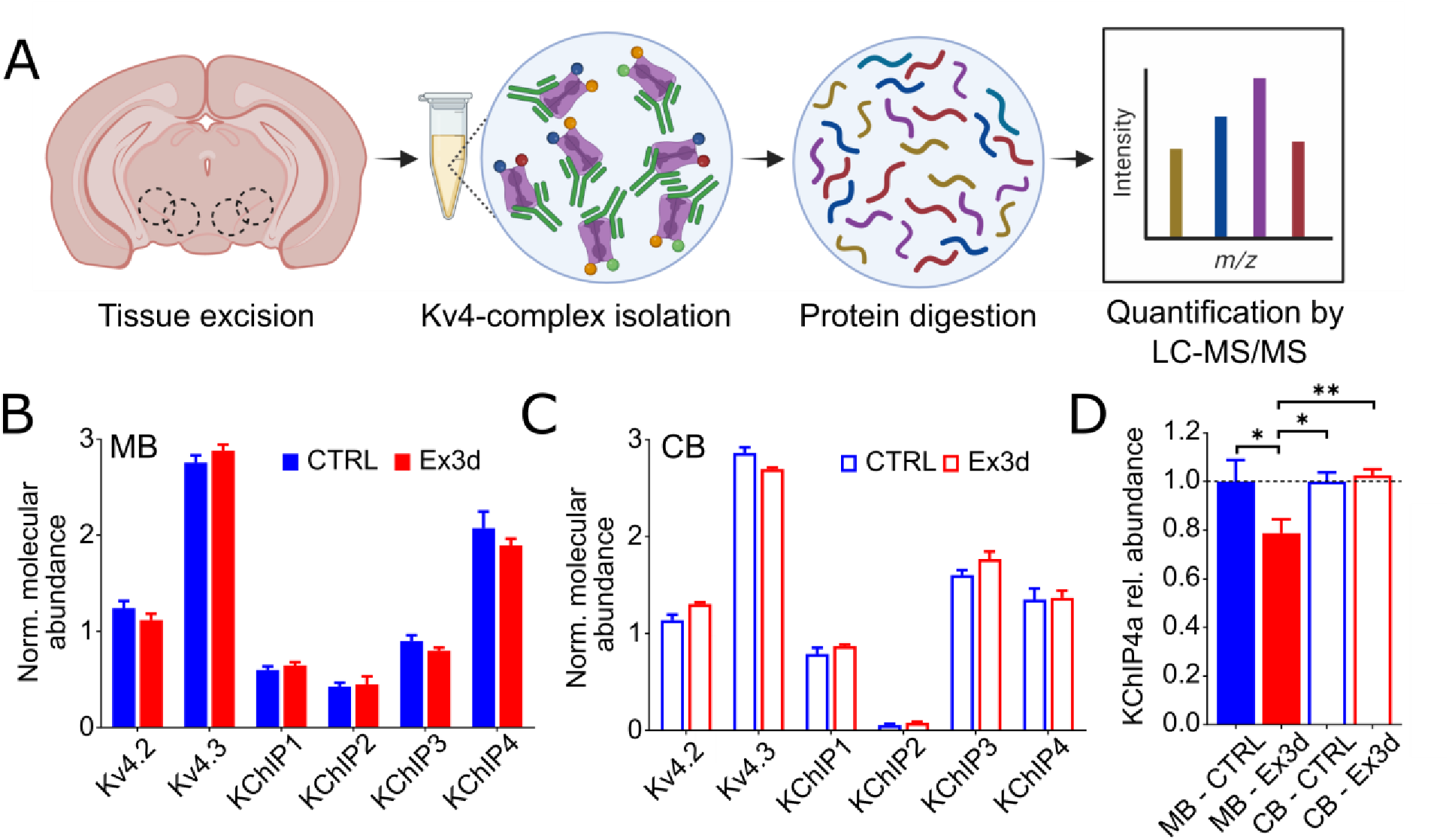
Ex3d mutation reduces KChIP4a protein expression in midbrain DA neurons without disrupting overall Kv4 channel subunit composition. **A:** Scheme of proteomic analysis of native Kv4 channel complexes. Tissue was excised from ventral midbrain and cerebellum, Kv4 channel complexes were affinity-isolated (antibodies in green, channels and associated proteins in other colors) and analyzed by liquid chromatography-tandem mass spectrometry (LC-MS/MS) following tryptic digest. **B, C:** Normalized molecular abundance determined for the indicated Kv4 complex constituents from ventral midbrain (MB; **B**) or from the cerebellum (CB, **C**) of both CTRL and Ex3d mice (see also Figure S2). Abundance was normalized to the tetrameric structure of the channel core. Note that Kv4.3 is the predominant pore-forming subunit (3/4 of each tetramer, on average), in either brain region, while KChIP4 is the prevailing auxiliary subunit in MB (2/4 of each tetramer, on average), but not in CB. **D:** Analysis of the molecular abundance of the KChIP4a protein (relative to CTRL) based on isoform-specific peptides. Note the selective reduction of KChIP4a abundance in the midbrain of Ex3d mice. N=12 experiments for Ex3d and CTRL. Error bars indicate standard error of the mean. *P < 0.05; **P < 0.01.

Next, we determined the relative abundance of KchIP4a in each tissue sample using MS-signals of isoform-specific peptides (see Methods). These analyses uncovered the selective reduction of KChIP4a protein in the midbrain samples from Ex3d mice (Figure 2C). The reduction was not complete (∼25%), likely because non-DA cells within the midbrain still express KChIP4a in Ex3d mice (Figure S1J), concordant with a targeted disruption of KChIP4a expression in midbrain DA neurons.

### KChIP4a regulates rebound delays and A-type currents in mesolimbic core DA neurons

To test whether KChIP4a was a molecular determinant of the long rebound delays observed in cNAcc-DA neurons ^4,6^, we performed *ex vivo* patch-clamp recordings of DA neurons from Ex3d and CTRL mice that projected to the NAcc core, NAcc lateral shell, or DLS, which were identified by retrograde axonal tracing and post-hoc IHC (Figure 3A, B and C). We found that the Ex3d mutation reduced the rebound delay in cNAcc-DA neurons by nearly 50 % (*P* = 0.016*; Figure 3D). Importantly, there was no effect on several membrane properties of DA neurons that projected to other striatal areas (Figure 3I and J), nor was there any observed additional effect in DA-cNAcc neurons (Figures 3D and S4). These results confirmed our hypothesis that KChIP4a is a main molecular determinant of the long rebound delays observed in DA-cNAcc neurons. Furthermore, this effect is highly selective, as no other variable was affected by the Ex3d mutation.

**Figure 3.**
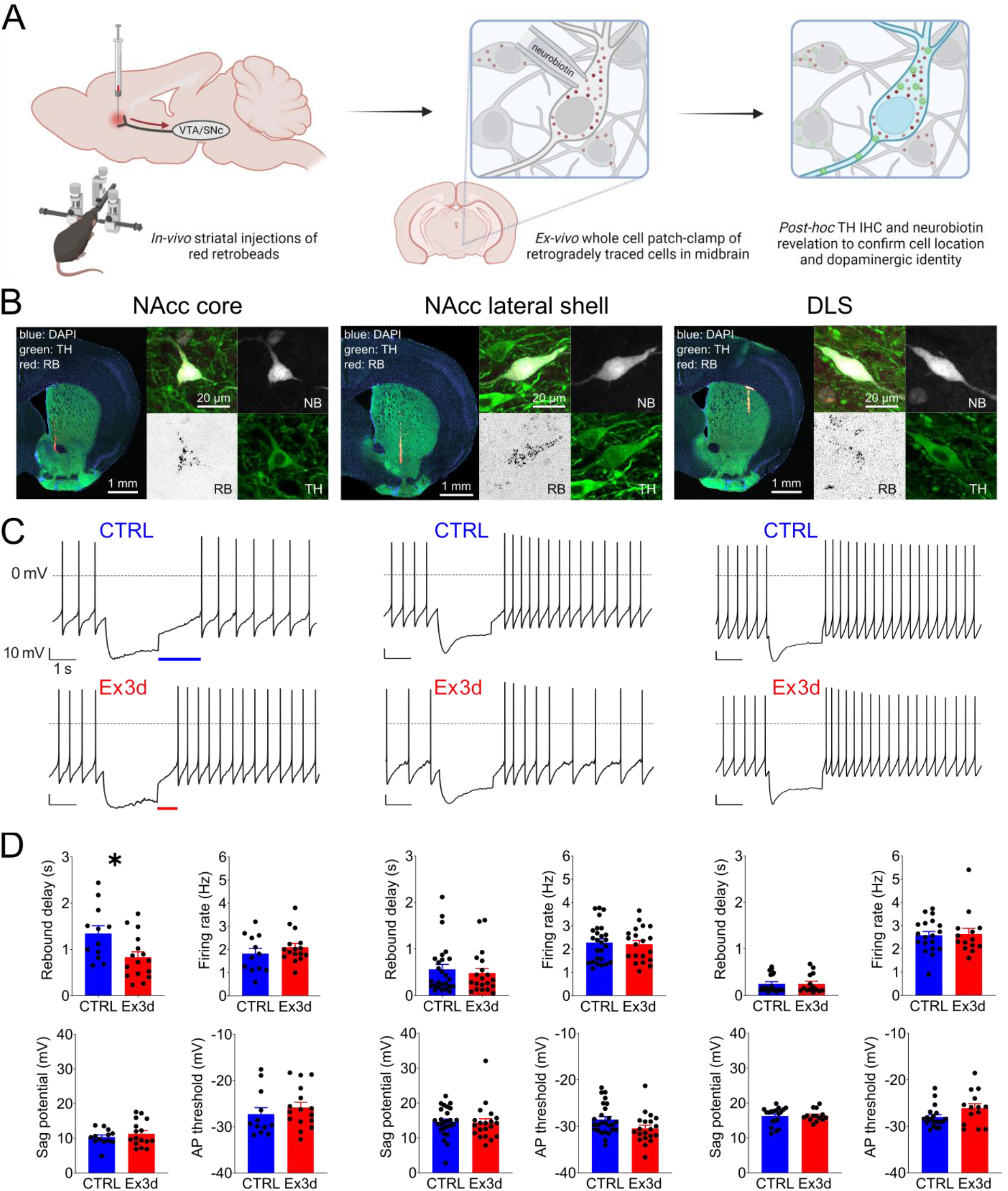
KChIP4a defines the longer rebound delay seen in NAcc core-projecting DA neurons. **A:** Schematic representation of the experimental timeline. Mice, CTRL and Ex3d, were injected with red retrobeads (RB) in different striatal areas, then retrogradely traced neurons in the midbrain were patched with neurobiotin-filled pipettes. After recordings, brain slices were processed for TH immunohistochemistry (IHC) and neurobiotin revelation, to confirm their DA phenotype and anatomical location, respectively. **B:** Representative micrographs of injections in the NAcc core, lateral shell and DLS, as well as of recorded, retrogradely-labeled, and TH-stained DA neurons from each of the projection pathways (See Figure S3 for the full mapping of injection and recording sites). **C**: Representative current-clamp recordings of DA neurons from both CTRL and Ex3d mice with confirmed projections to the NAcc core (N=16/4 for Ex3d and N=12/4 for CTRL), lateral shell (N=20/4 for Ex3d and N=27/4 for CTRL) or DLS (N=14/3 for Ex3d and N=19/3 for CTRL), in order (left to right), in response to a -80 mV hyperpolarizing pulse. Note the shorter rebound delay in the NAcc core-projecting example from an Ex3d mouse in relation to CTRL. **D:** Biophysical properties of projection-identified DA neurons from CTRL and Ex3d mice. The only significant difference observed across genotypes and projection targets was a significant shortening of rebound delay in DA-cNAcc neurons of Ex3d mice in relation to CTRL. See Figure S4 for more comparisons. AP = action potential. Error bars indicate SEM. *P < 0.05.

To confirm the biophysical underpinning of the difference in rebound delays, we performed voltage clamp recordings of A-type currents in DA-cNAcc neurons of Ex3d and CTRL mice. We found that KChIP4a deletion had a dual effect on I_A_ kinetics, speeding up both open-state inactivation (*P* = 0.002**; Figure 4A) and recovery from inactivation (interval effect *F*_3, 48_ = 110.2, *P* < 0.0001***; interaction of interval and genotype effect, *F*_3, 48_ = 5.321, *P* = 0.003**; Figure 4B). These results are in line with previous descriptions of KChIP4a’s effect on A-type currents in heterologous systems ^7,9^.

**Figure 4.**
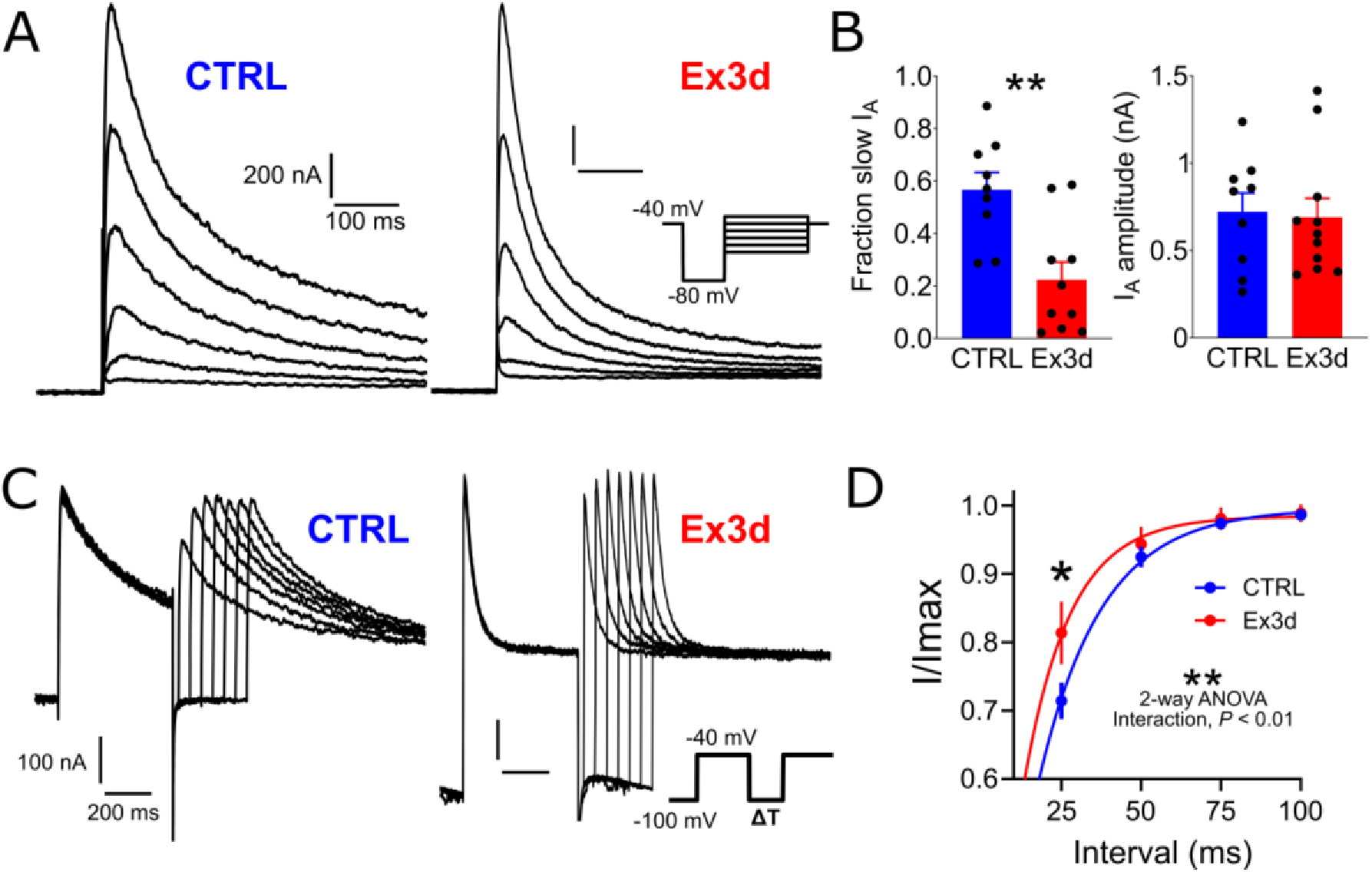
KChIP4a controls A-type current inactivation and recovery in NAcc core-projecting DA neurons. **A:** Representative recordings of A-type currents (IA) from CTRL and Ex3d DA-cNAcc neurons, elicited with successively larger depolarization pulses from -80 mV. **B:** In relation to controls, A-type currents recorded from neurons of Ex3d mice had a significantly lower fraction of slow inactivation, i.e., faster inactivation kinetics, but showed no difference in peak amplitude. **C:** Recovery from inactivation was tested by applying two 60 mV pulses from -100 mV separated by increasing intervals. **D:** Compared to CTRL, Ex3d cells showed faster recovery from inactivation. (N=9/ for Ex3d and N=9/5 for CTRL). Error bars indicate SEM. *P < 0.05.

### Simulation of KChIP4a deletion reveals a shift in inhibitory integration

Given that the dynamics of *in vivo* subthreshold input integration in DA neurons have only been inferred from intracellular recordings in anesthetized animals ^23–25^, the ultimate consequence of KChIP4a removal on behaviorally-relevant neural activity could depend on several unknown parameters, such as pre-synaptic neurotransmitter release dynamics. In order to form a specific hypothesis for the behavioral consequences of KChIP4a deletion in DA neurons, we needed to infer how the potential effects on input processing varied across different conditions of synaptic connectivity and temporal integration. To do this, we used biophysically realistic computational models of DA-cNAcc neurons (Figure 5), tailored to reflect their empirically defined membrane properties ^26^.

**Figure 5.**
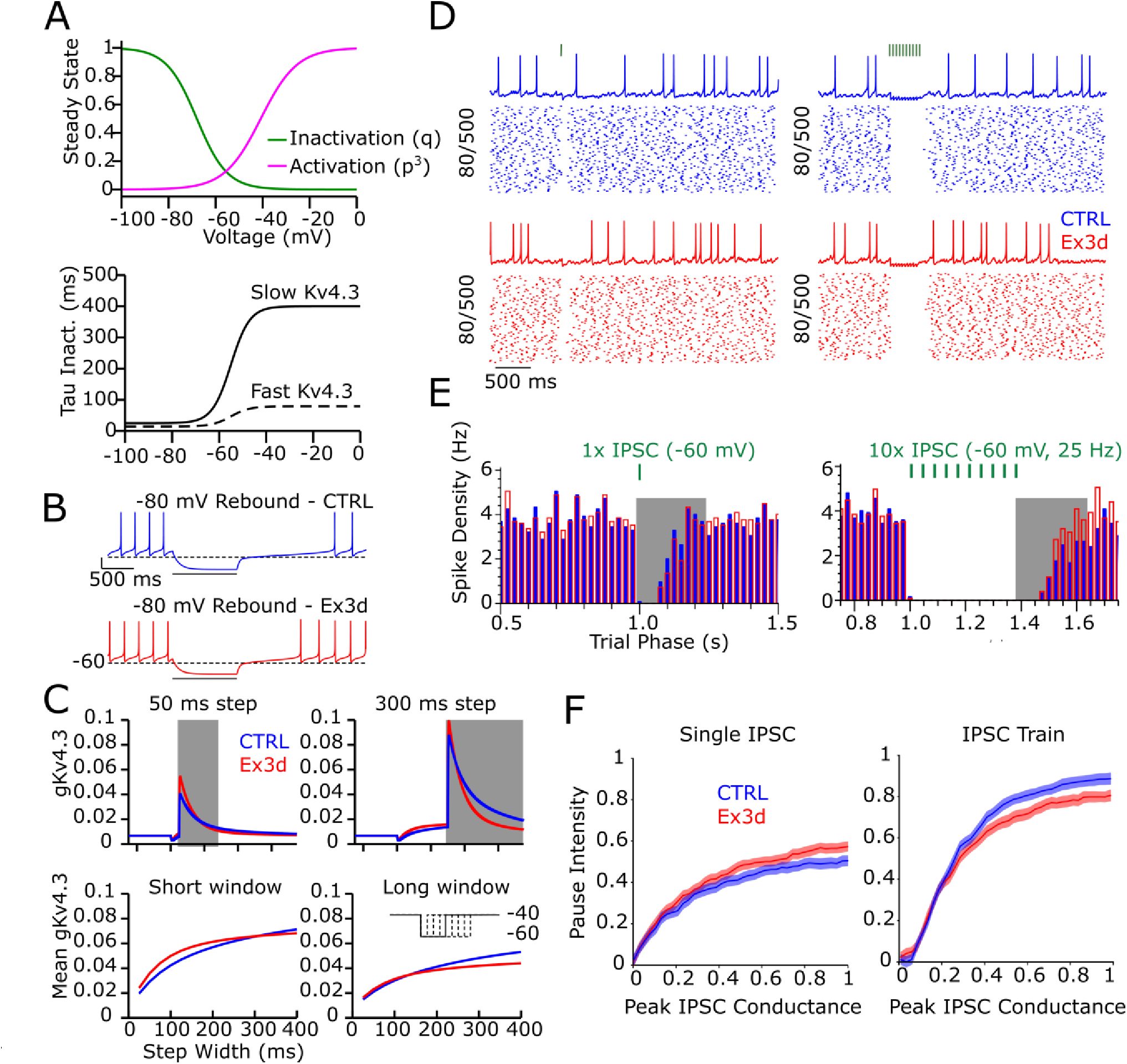
Modelling of the biophysical effects of KChIP4a deletion in DA neurons predicts altered *in vivo* firing pauses after inhibition in Ex3d mice. A: Model calibration of Kv4.3 models, including activation and inactivation curves for all Kv4.3 subunits (top) and time constant of the inactivation gating variable for KChIP4a (solid) and fast Kv4.3 subunits (dashed) (bottom). B: Model rebound response from 1s step to -80 mV for CTRL subunit distribution (blue) and Ex3d subunit distribution (red). C: Short hyper-polarizations predicted to preferentially recover fast Kv4.3. Top panels show total Kv4.3 conductance of CTRL (blue) and Ex3d (red) at -45 mV following a 50 ms (left) and 300 ms (right) step to -60 mV. Left bottom panel shows the time-averaged Kv4.3 conductance during the 100 ms post rebound window (gray box in panel above) for CTRL and Ex3d following -60 mV voltages steps ranging from 25 and 400 ms, while right bottom panel shows the same during the 250 ms post rebound window (gray box in above panel). Inset represents command voltage protocol. D: Representative traces and raster plots for CTRL (blue) and Ex3d (red) model neurons in a simulated 3.7 Hz balanced state perturbed by either single (left) or multiple (10x IPSC at 25hz, right) identical simultaneous GABAA (ECl- = -60 mV) inputs. E: Spike density histogram of trials in D (aligned to above). Bins are 25 ms intervals averaged over 500 total trials. Density of CTRL (solid blue) overlayed with Ex3d (red box). Grey overlay indicates a 250 ms rebound interval. F: Pause intensity (decrease in spike density) in rebound interval as IPSC amplitude (gGABAA) is varied over the fixed balanced state for single IPSCs (left) or a train of 10 IPSCs at 25 Hz (right). Intensity is normalized to that of unperturbed balanced state. Error bands correspond to the standard deviation of the mean.

To simulate the effects of binding of different KChIPs, we modelled the Kv4.3 with a Hodgkin-Huxley style ^27^ model with two independent additive inactivation states (Eq, N; slow and fast, respectively). The slow gate is associated with the conductance of channels modulated by the KChIP4a subunit, whereas the fast gate is associated with the conductance of all other “fast” subunits ^7,9^. The steady state values for both inactivation and activation (Figure 5A), along with the kinetics of activation, were adapted from Tarfa et al (2017), and in the absence of population-specific data, steady-state parameters were held constant across subpopulations (Table S1). The kinetics of the fast and slow inactivation gates are given in (Figure 5A) and were taken from the current clamp data presented in Figure 4. Representative cells for both CTRL and Ex3D were created using the mean ratios and peak currents from Figure 4B for those respective data sets. Consistent with *ex vivo* rebound data in Figure 3C, the CTRL model cell has a rebound delay of 1.9 second when released from a 1 second step to -80 mV, while the Ex3D model cell has a rebound delay of 0.97 second under the same conditions (Figure 5B). Additionally, consistent with recorded data, the basal firing rate (3.4 in CTRL and 3.3 in Ex3D) was not significantly affected.

Next, we explored how these different Kv4.3 currents could affect synaptic integration. *In vivo,* hyperpolarization rebound delays are concurrent with a barrage of both excitatory and inhibitory inputs in the balanced state. Thus, the duration of post-inhibitory pauses is not necessarily determined by the length of the biophysically-determined rebound delay. Rather, it is shaped by the integration of this rebound with concomitant synaptic input. Kv4.3 channels that are de-inactivated during hyperpolarization produce a large outward current upon release from hyperpolarization with a conductance that increases rapidly during the subsequent depolarization. The fast activation of this channel, on the order of 1 ms ^6^, creates a restorative counterbalance not only to intrinsic inward currents, but glutamatergic EPSPs as well.

Due to the differences in the timescales of both inactivation and recovery from inactivation between the two subunits, we hypothesized that the CTRL and Ex3D cells would respond differently to rebound following short and shallow hyperpolarizing steps, which would mostly de-inactivate “fast” Kv4 channels (without KChIP4a), versus longer hyperpolarizing steps, which would fully de-inactivate channels with all subunits. We tested this hypothesis by comparing the total Kv4.3 conductance of the Ex3d and CTRL models in response to a simulated voltage clamp experiment holding the cell at a typical subthreshold voltage value of -40 mV with variable length pulses to a typical GABA_A_ reversal potential of -60 mV. For short (25-200 ms) hyperpolarizing steps, such as those that would be created by a brief inhibitory input, the Ex3d cells produced 25 % larger initial Kv4.3 currents that decayed below that of the CTRL after about 100 ms (Figure 5C). For longer (>200 ms) hyperpolarizing steps consistent with a train of inhibitory inputs, the initial currents were roughly equivalent, but the faster subunits in Ex3d inactivate more quickly, leaving a larger conductance in the CTRL over longer timescales.

Comparing the mean open fraction of Kv4.3 channels of the CTRL and Ex3D models over post-inhibitory windows of 100 ms and 250 ms (Figure 5C) reveals that over short timescales Ex3D cells are more efficient at producing quick surges in Kv4.3 conductance, while CTRL cells are more efficient at producing long lasting Kv4.3 conductance from longer pauses.

To probe the potential consequences of this finding to the processing of behaviorally relevant information, we tested how CTRL and Ex3d cells would respond in a condition that mimicked as closely as possible the synaptic environment in an intact mouse (a “simulated *in vivo*” model). We did this by applying simultaneous Poisson process trains of excitatory and inhibitory postsynaptic potentials (EPSPs and IPSPs, respectively), to create a balanced state similar to what has been experimentally observed in DA neurons *in vivo* ^23^, *ex vivo* using dynamic clamp ^28^, and in computational models ^29^. We then applied phasic inhibitory pulses by introducing temporally correlated GABA_A_ inputs on top of the existing balanced state inputs (Figure 5D). For both single pulse inhibition and a short train of inhibition (10 pulses at 25 Hz; Figure 5D), examination of the individual traces is not sufficient to reveal subtle differences in population responses. Repeated presentations of the stimulus to a single noisy neuron, analogous to trial-based experimental approaches for studying DA neuron correlates in reinforcement learning ^1,13^, are required to characterize the differences in the response of CTRL and Ex3d cells. Consistent with the larger mean Kv4 conductance evoked during short time windows following brief hyperpolarization pulses (Figure 5C), Ex3D model neurons had significantly larger pause intensities (effectiveness to decrease firing) immediately following single pulse inhibition relative to CTRL (Figure 5E). Also consistent with the smaller Kv4 conductance evoked during longer time windows following more long-lasting hyperpolarization (Figure 5C), Ex3D models exhibited significantly lower pause intensities immediately following long inhibitions (Figure 5E). These differences were found over a wide range of input strengths, with the model Ex3D cell saturating at a 13 % higher pause intensity following pauses (Figure 5F). In contrast, there were 53 % less pauses spikes immediately following the 360 ms pulse of 25 Hz inhibitory input. Taken together, this suggests that Ex3D cells have an elevated gain in translating short, temporally correlated inhibitory signals into pauses at a network level. This comes at the cost of a reduced gain in translating trains of inhibitory input into more sustained pauses. Taken together, our *in silico* exploration of the effects of KChIP4a deletion in DA-cNAcc neurons predicted that Ex3d mice could show more effective hyperpolarization-induced pausing *in vivo*, especially regarding transient synaptic events, which should consequently enhance learning from negative RPEs.

### KChIP4a deletion in DA neurons selectively enhances learning from negative RPEs

While our modelling-inspired hypothesis was very straightforward, we subjected Ex3d mice and CTRL to a battery of DA-sensitive tasks to test how the deletion of KChIP4a in DA neurons could have more general effects on behavior. This included open field exploration (Figure 6A), hole board exploration (Figure 6B), novel object preference (Figure 6C), and spontaneous alternation in the plus maze (Figure 6D). There was no difference between the genotypes in any of these tasks, indicating that deleting KChIP4a in DA neurons did not impact locomotion, anxiety, spontaneous exploratory behavior, novelty preference and learning, short-term memory, nor working memory. This series of tests demonstrated that KChIP4a deletion in DA neurons does not produce a widespread effect on DA-dependent tasks.

**Figure 6.**
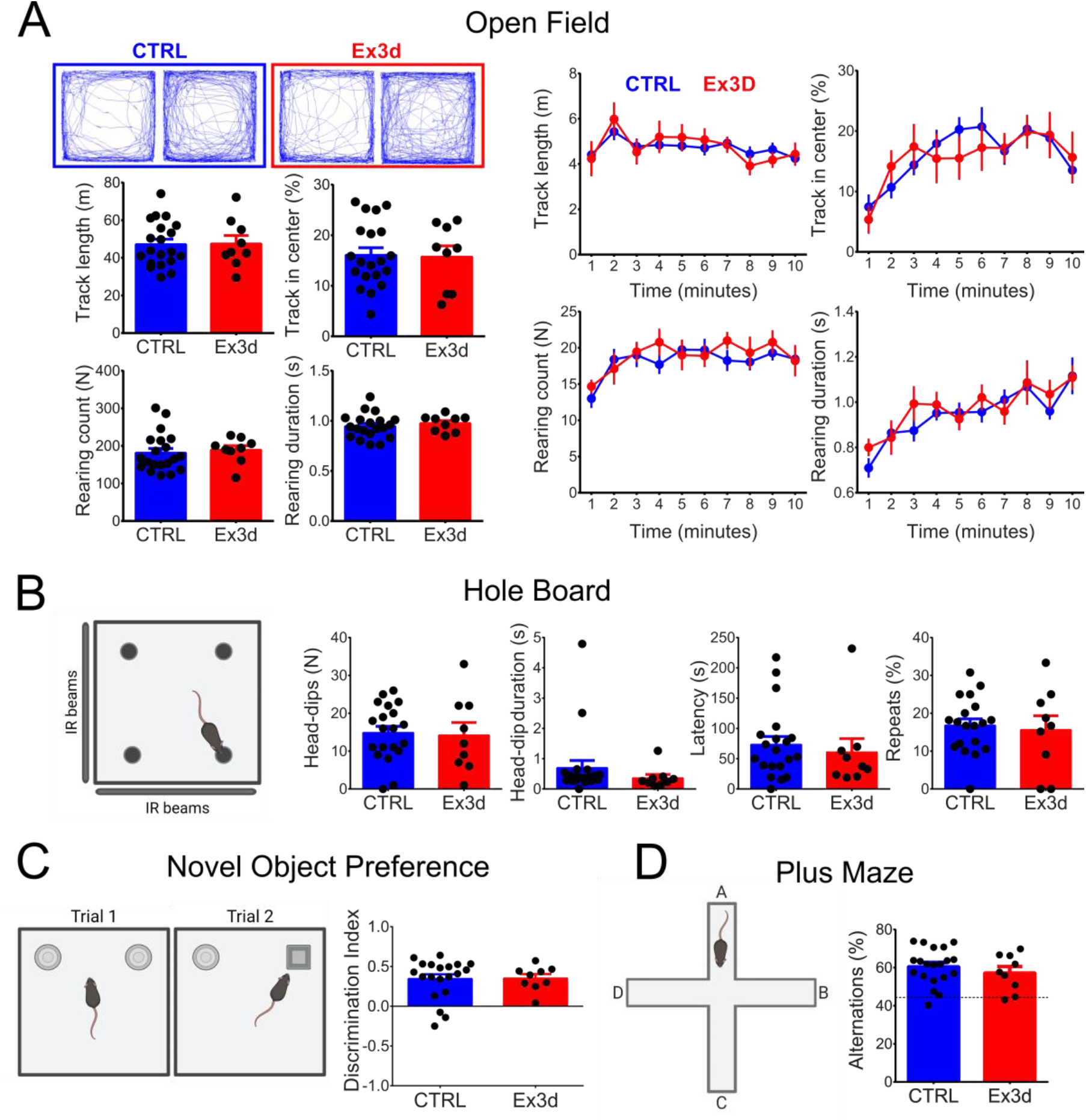
DA neuron-specific KChIP4a deletion does not affect several DA-dependent behaviors. **A:** Session-based results of the open field test, including representative tracks from mice of both genotypes, averages of the total track length, proportion of track length in the center of the open field, number, and duration of rearings during the whole 10 minutes of the task and averaged across every minute of the 10 minute task. **B:** Hole board test results, including total number of head dips, total summed head-dip durations, latency to first dip and of repeat head-dips. **C:** Cartoon representation of the novel object preference test, and the results of the discrimination index (positive values indicate preference for the novel object in trial 2). **D:** Plus maze test results; spontaneous alternations were quantified as the proportion of spontaneous alternations relative to the total possible alternations. Note the absence of a significant genotype difference for all variables in all tests (unpaired T tests and two-way repeated measures ANOVA, P > 0.05). N=9 for Ex3d and N=20 for CTRL. Error bars indicate SEM.

To test the model-inspired hypothesis that learning from negative RPEs should be increased by KChIP4a deletion, we trained mice on a reinforcement learning task where they learned that an auditory cue signaled the availability of sugar water reward (acquisition), followed by sessions where reward was omitted (extinction). We did not observe a significant difference between genotypes on any behavioral variable during acquisition (Figure 7A-D). However, we found that, as predicted by our biophysical modelling, Ex3d and CTRL mice differed dramatically in extinction learning. The Ex3d group displayed a significantly accelerated reduction in the time they spent in the reward port during cue presentation (interaction of genotype and session progression effect, *F*_5, 135_ = 3.192, *P* = 0.001**; Figure 6A) and a faster increase in response latency (interaction effect, *F*_5, 135_ = 2.615, *P* = 0.027; Figure 6B) across extinction sessions.

**Figure 7.**
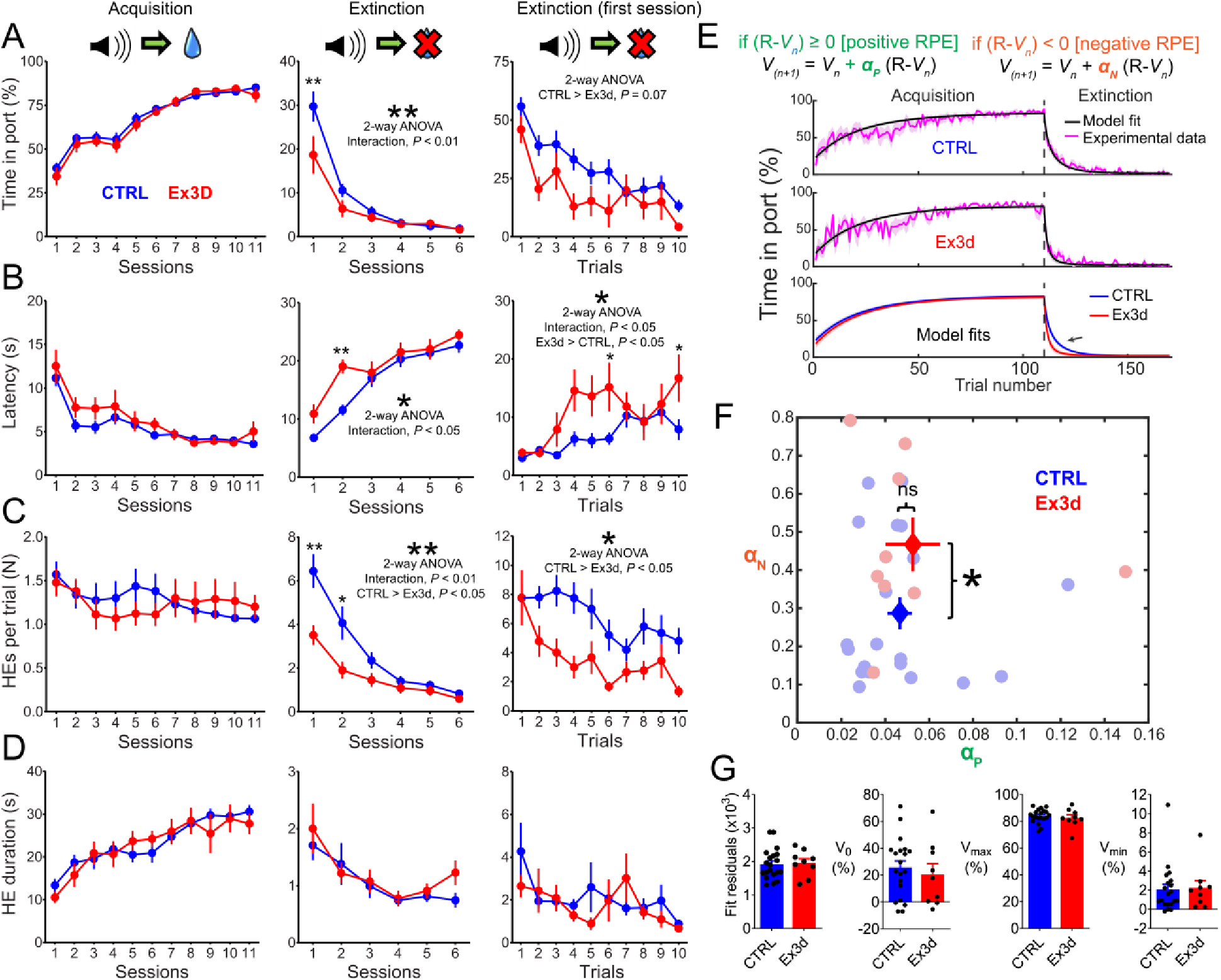
DA neuron-specific KChIP4a deletion selectively improves learning from negative RPEs. **A:** Time spent in port during CS presentation on a session-by-session basis during acquisition and extinction, as well as on a trial-by-trial basis on the first extinction session. Note the faster response extinction for Ex3d mice in relation to controls, including the significant pair-wise difference during the first extinction session, and the strong trend towards faster trial-by-trial extinction in the first extinction session. **B:** Latency to respond during CS presentation for all phases of the reinforcement learning task. This measure increased faster during extinction in Ex3d mice. Importantly, there was also faster trial-by-trial learning in the first extinction session. **C:** Number of head entries into the reward port during CS presentation. Note the faster response extinction for Ex3d mice in relation to CTRL, as well as a faster trial-by-trial extinction in the first session of extinction in Ex3d mice. **D:** Duration of head entries into the reward port during CS presentation. Note the lack of a genotype effect during all phases of acquisition and extinction. Furthermore, for all measured variables, there was no significant difference between genotypes during acquisition or in the first trial of the first session of extinction. **E:** Fitting of a Rescorla-Wagner model to the trial-by-trial time in port data. Note that the model tracks the behavioral data with relatively good accuracy in both groups, and that the model fits overlap almost completely during acquisition, but diverge in extinction, reflecting the faster extinction learning kinetics of the Ex3d mice in relation to controls (black arrow). **F:** Best fit αP and αN values for Ex3d (red) and CTRL (blue) groups. Circles represent individual values. Crosses indicate the mean ± SEM of each group. Note the selective increase in αN (learning from negative RPEs) in Ex3d in relation to littermate controls. **G:** V0, Vmin, Vmax, and best fit residuals for Ex3d and CTRL groups. No significant difference was observed between the compared groups. N=9 for Ex3d and N=20 for CTRL. Error bars indicate SEM. *P < 0.05; **P < 0.01.

To verify whether we were observing a direct effect on learning (which would be compatible with an impact on RPE signaling) or a reduction of responding in the absence of direct reinforcement (more compatible with an impact on motivational processes), we analyzed the trial-by-trial behavior measures in the first extinction session. If the Ex3d mice already displayed a reduced time in port on the first extinction trial (when the mice were first confronted with an unexpected reward omission), then the genotype effect would be more likely attributable to decreased motivation ^30^. However, we found that Ex3d and CTRL mice were statistically indistinguishable in their responses during the first extinction trial (Figure 7A-D). Only in later trials of the first extinction session, genotype differences emerged in response latency (interaction effect, *F*_9, 243_ = 2.01, *P* = 0.031*; genotype effect, Ex3d > DAT-Cre KI, *F*_1, 27_ = 7.51, *P* = 0.01*; Figure 7B) and in the number of head entries (genotype effect, CTRL > Ex3d, *F*_1, 27_ = 5.98, *P* = 0.021*; Figure 7C). We also identified a strong trend for a gradual divergence between genotypes, starting on the second trial, for the time in port metric (genotype effect, CTRL > Ex3d, *F*_1, 27_ = 3.51, *P* = 0.07; Figure 7A). This confirms that the Ex3d effect developed progressively as the mice learned that the CS no longer predicted US availability, strongly suggesting a selective effect on learning from expected reward omission, i.e., negative RPEs.

To formally test the hypothesis that only negative RPEs were affected, we fitted the trial-by-trial time in port values for both Ex3d and CTRL groups with a Rescorla-Wagner model which assumed different learning rates for positive (α_P_) and negative (α_N_) RPE-based learning ^31–33^. This model successfully recapitulated the behavioral data, including the different learning kinetics during extinction (Figure 7E). Comparisons of the learning rates for the best fits showed that indeed the Ex3d group had significantly higher α_N_ values, i.e., faster learning from negative RPEs (*P* < 0.05), without any change in α_P_ (Figure 7F). Importantly, empirically determined model parameters (*V*_0_, *V*_max_ and *V*_min_), as well as goodness of fit, were similar for both genotypes (Figure 7G). These results support the interpretation that the effect of deleting KChIP4a in DA neurons can be explained by a selective increase in learning from negative RPEs.

## Discussion

In this study, we disrupted a specific genetic mechanism – the alternative splicing of the Kv4 channel regulatory subunit KChIP4a – in DA neurons and tracked the selective effects of this perturbation across multiple scales ranging from Kv4 channel complex composition and gating, cellular membrane properties, information integration, to behavior. We achieved this by generating a transgenic mouse line where KChIP4a is selectively removed from DA neurons, confirming the precision of our molecular perturbation at the level of mRNA and Kv4 channel complex composition using FISH and high-resolution proteomics.

We then showed that this deletion reduced rebound delays in DA-cNAcc neurons, while sparing both other physiological variables within these neurons and all properties of other DA subtypes, namely DA-lNAcc and DA-DLS neurons. This demonstrated that KChIP4a is a molecular determinant of the previously described variability in rebound delays between different subpopulation of DA neurons ^4,6^. The subpopulation specificity of this phenotype is likely due to the intrinsic differences in the expression KChIP4a between the cell groups – DA-cNAcc neurons may just express more KChIP4a and are thus more sensitive to its deletion. It could also be determined by differences in relative HCN currents, which typically oppose the effects of A-type currents by accelerating rebound delays. Cells that barely have hyperpolarization-activated depolarizing currents (a.k.a. sag), like DA-cNAcc neurons, would have only a limited range for compensating the effects of KChIP4a deletion, while cells with more sag could potentially adjust HCN channel expression to match the changes in Kv4 channel properties. It is also possible, or even likely, that both mechanisms are involved in determining the final phenotype. The biophysical selectivity of the phenotype could be attributable to the fact that Kv4 channels in atypical DA VTA neurons are mostly inactivated at typical neuronal resting potentials and require hyperpolarization to recover from inactivation ^8^, i.e. they contribute little to the cell’s electrophysiological dynamics unless there is hyperpolarizing inhibition. This is in contrast to the pacemaker control function of Kv4 channels in conventional DA SN and DA VTA neurons ^5,34^.

We further used biophysical computational models of DA-cNAcc neurons to simulate the effects of KChIP4a deletion across a wide variety of conditions. We demonstrate that the shift in inactivation kinetics produced by the removal of KChIP4a is enough to shorten rebound delays, supporting the conclusion that KChIP4a is the biophysical underpinning of long rebounds in atypical DA-cNAcc neurons. Effectively, relative to CTRLs, Ex3d DA-cNAcc neurons have more Kv4.3 channels available for activation very shortly after, but less so after a longer time period (as they inactivate faster and therefore lead to lower general conductance).

To infer the *in vivo* functional consequences of such a change, we used Poisson-distributed IPSP and EPSP trains to simulate an *in vivo*-like inhibition-excitation balance and probed how this biophysical switch could affect information integration. We found that barrages of IPSPs, which supposedly underlie negative RPE integration in DA neurons ^13,31^, tested across several combinations of IPSC number amplitude, led to different firing pause durations in Ex3d models versus controls. Interestingly, the *in silico* removal of the slow inactivation component associated with KChIP4a modulation made pauses longer when inhibitory inputs were short and shorter when they were long, similar to the observed effect on Kv4 conductance, and therefore systematically shifting the integration of GABAergic stimuli.

Finally, we show mice with the Ex3d mutation induced a highly specific change in learning from negative RPEs, with several other DA-dependent behaviors being spared. This deletion had a circumscribed effect on the learning rate during extinction of a conditioned response. It did not affect acquisition or other behavioral processes like locomotion and working memory. Moreover, this effect on extinction learning was circumscribed to changes in the rate with which mice initiated reward-seeking behaviors, without affecting disengagement dynamics ^35^. Fitting the behavior data to a reinforcement learning model confirmed that this phenotype is compatible with a selective increase in negative RPE learning rates ^36^.

Several studies have shown that pause duration and firing rate decreases in DA neurons, as well as reductions in DA release or DA axon Ca^2+^ transients in the ventral striatum, are directly correlated with negative RPE signaling ^14,31,37,38^. Our results show that DA-cNAcc neuron inhibition, and subsequent negative RPE signaling, are under tight biophysical control by a unique K^+^ channel modulator. The most straightforward interpretation of our findings, in line with the aforementioned literature, is that the removal of KChIP4a from these cells increases pause durations following negative RPE-related IPSCs. This explicitly predicts that normal signaling of negative RPEs involves surges in synaptic inhibitory currents that are somewhat small, infrequent, or both, as these would be amplified by KChIP4a deletion.

Interestingly, some recent studies, using precise optogenetic inhibition of DA neurons during reinforcement learning tasks, have suggested that negative RPEs are more effectively signaled by *multiple short pauses* rather than by *single long pauses* in DA neuron firing ^16,18,39^. While this in some ways is in line with our prediction that endogenous negative RPEs are signaled via short inhibitory bouts, it also could suggest the alternative, albeit less likely, possibility that KChIP4a deletion increases learning from negative RPEs by reducing pause durations or even by breaking apart long firing interludes into multiple short breaks. Future work should directly test these different explanations.

There are interesting possibilities as to how altered inhibition integration in DA-cNAcc neurons is translated into stronger negative RPE-based learning at a population level. It was recently shown that VTA DA neuron ensembles encode RPEs in a distributed manner, i.e. individual DA neurons encode RPEs with different biases, so that as a population they can fully represent the distribution of reward availability contingencies ^40^. Individual DA neurons fired as if operating with different positive and negative RPE learning rates – α_P_ and α_N_ – and therefore were differentially affected by rewarding or disappointing events. The mechanisms that determine this heterogeneity were, however, not determined. We speculate that, by specifically controlling α_N_, KChIP4a expression could be important for setting this positive/negative RPE signaling bias in each cell. Hence, removing KChIP4a from DA neurons may have shifted the population-level encoding of RPEs towards a more “pessimistic” bias, leading to faster extinction.

Kv4 channels, and their regulation by auxiliary subunits, are also key modulators of synaptic integration in other, non-dopaminergic, cell types. For example, in olfactory bulb mitral cells, fast Kv4 currents selectively attenuate AMPA-, but not NMDA-mediated EPSCs ^41^, and in CA1 pyramidal neurons, Kv4 currents control the amplitude and duration of EPSCs, the backpropagation of dendritic action potentials, and the threshold for LTP induction ^42,43^. Our results add to this literature by demonstrating, to our knowledge for the first time, that slow Kv4.3-KChIP4a currents can act as relatively selective regulators of hyperpolarizing inhibitory inputs, a feature that may be present in other cell types.

Two remarkable aspects to our findings are 1) that the deletion of a single splice variant produced such a highly penetrant phenotype across multiple scales of organization, and 2) that these effects were strikingly selective at all levels of analyses. While much recent work argues that neuronal ion channel expression is typically degenerate, i.e., the function of different conductances overlap to ensure functional robustness ^44^, our findings would suggest that this is not the case for KChIP4a in DA-NAcc neurons. This may be because the unique properties of this subunit are particularly vulnerable to loss of function mutations, leading to a high selection pressure for feature conservation. This would explain why the structure of KChIP4a is more similar to the earliest known ancestral KChIP protein than other members of this family ^7,45^. DA-NAcc neurons, due to their own ion channel profile, seem to be reliant on KChIP4a as a tuner of inhibitory integration. Other cell types, neuronal or not, could also be similarly reliant on KChIP4a. As a general principle, perhaps some channels, or their modulatory subunits, cannot be functionally compensated by degenerate mechanisms, and are therefore critical nodes for determining cellular function and disfunction.

Our findings may also be relevant for the understanding of human disease. Recent studies have demonstrated that surviving nigral DA neurons of parkinsonian patients and animal models of parkinsonism have increased KCNIP4 expression ^46,47^. While these studies did not differentiate between different KCNIP4 isoforms, it could be that the differential expression of KChIP4a plays a key role in determining DA neuron resilience in Parkinson’s disease, especially since nigral DA neurons in animal models of parkinsonism show reductions in Kv4 currents ^48,49^.

Furthermore, polymorphisms in the KCNIP4 gene have been linked to multiple mental illnesses, including substance use disorder ^50–53^, attention-deficit-hyperactivity disorder ^54–56^, depression ^57,58^, bipolar disorder ^59,60^, autism ^61^, and several personality disorders ^54^. Interestingly, most of these polymorphisms are in intronic regions, suggesting that regulatory changes in KCNIP4 gene expression, including differences in splicing cis-regulatory elements, may play a key role in these associations ^62^. Furthermore, the diseases most frequently associated with KCNIP4 variations – substance use disorder, ADHD, depression, and bipolar disorder – have been directly linked to altered learning from negative RPEs ^63–66^. Our findings raise the intriguing hypothesis that KCNIP4 polymorphisms may increase predisposition to certain mental illnesses by, changing the regulation of KChIP4a expression and subsequently affecting negative RPE-based learning. Given the relative lack of degeneracy for KChIP4a function, this subunit may be particularly important for defining vulnerability to disease.

In summary, we have identified KChIP4a as a selective molecular modulator of inhibitory integration in a subset of DA neurons and, consequently, of learning from negative RPEs, singling out this subunit as a likely homeostatic controller of adaptive behaviors. Our findings demonstrate the existence of a highly specialized gene-to-behavior mechanistic chain built within the midbrain DA system, furthering our understanding of how the genetically determined biophysical diversity of DA neurons may be critical for the execution of specific computational processes.

## Acknowledgments

The authors would like to thank Beatrice Fischer, Jasmine Sonntag, and Felicia Müller-Braun for their exceptional technical support. This research was supported by DFG CRC 1080 (A11) to JR and by NIH grant R01DA041705 to CCC and JR. KMC received a one-year fellowship and financial support from the Max Planck Society (International Max Planck Research School for Neural Circuits – Max Planck Institute for Brain Research).

## Author contributions

KMC and JR designed the overarching research project, interpreted all results, and wrote the manuscript. JR supervised the research project. JR and CCC acquired financial support for the research project. KMC wrote the first draft of the manuscript, planned, performed, and analyzed results from behavioral experiments and behavioral modelling, and compiled data from all experiments. NH and JR planned, performed, and analyzed results from the *ex vivo* electrophysiology experiments. TM and DS planned, performed, and analyzed results from the combined FISH/IHC assays. JS and BF planned, performed, and analyzed results from the proteomic experiments. CK and CCC developed and implemented the biophysical modelling of DA neurons.

## Methods

### Animal protocols

All animal procedures described in this study were approved by the German Regierungspräsidium Gießen and Darmstadt (license numbers: V54-19c 20/15 –FU/1100 & FU1203). Mice were bred and housed until 8 weeks of age at MFD diagnostics (Mainz, Germany). Mice were then maintained under a 12 h dark/light cycle and housed in groups of two to four, with food (R/M-keeping, Ssniff, Germany) and tap water available *ad libitum*, except when they underwent water restriction. Nesting material and a red acrylic glass shelter (mouse house, Tecniplast, Germany) were used as enrichment. Both males and females were used for all experiments - we did not observe any evident effect of sex in any of the experiments, and therefore pooled together data from both sexes. For every single experimental procedure in this study, the experimenter was blind to the genotype of the mice and the order in which mice were tested was pseudo-randomized by another team member that was not directly involved in the execution of the experiment.

### The DA-neuron selective KChIP4a <u>ex</u>on <u>3 d</u>eletion (Ex3d) mouse line

The mouse KChIP4 gene, KCNIP4, is located on chromosome 5 (B3, 48.39 – 49.52 Mb), is 1135 kb in length and has 14 exons, with ATG translation initiation codons in exons 1, 2, 3, 5, and 8, and the STOP codon located in exon 14 ^67^. KChIP4a (a.k.a. KChIP4.4) is a 229 amino acid protein that corresponds to the alternative splicing of exons 3 through 8b of KChIP4, with exon 3 coding for the KID ^9,68^.

A new transgenic mouse line was developed for assessing the specific function of the KChIP4a KID using the Cre-lox system. Exon 3 of the KCNIP4 gene (which encodes the KID) was flanked with loxP sites using a vector designed to display two homology regions in a C57BL/6N genetic background, including a short homology region of 1795 bp and a long region of 5858 bp. This vector was electroporated into C57BL/6N embryonic stem cells and screened clones were used for the generation of mice with homozygotic floxing of KCNIP4 exon 3 (*KChIP4-Ex3^lox/lox^*). These mice were then crossed with a line with heterozygous DAT-Cre knock-in (*DAT-Cre^+/-^*or *DAT-Cre KI*) mice. The offspring of this crossing were then interbred, leading to the production of mice where exon 3 of the KCNIP4 gene was selectively excised from DAT-expressing neurons (Ex3d; *DAT-Cre^+/-^*/ *KChIP4-Ex3^flox/flox^*) and DAT-Cre KI control littermates (CTRL; *DAT-Cre^+/-^ / KChIP4-Ex^WT/WT^*).

Genetic identity of all KChIP4-Ex3^flox/flox^ mice was confirmed with polymerase chain reaction (PCR) genotyping throughout the breeding process. For detecting the conditional deletion allele, the following primers were used:

5-TAG TTA TGA CAA GAC AGG AGC TAG TAC CAC TAA GC-3

and

5-GAA CTG GAC TGA AGC AAA ACA AAA CAC G-3.

These primers target the flanking region of the FRT site and the PCR product lengths were 385 bp for the WT allele and 477 bp for the conditional deletion allele. The PCR protocols were run with 1 cycle at 94 °C for 120s (denaturing), 35 cycles of 94 °C for 30s (denaturing), 65 °C for 30s (annealing) and 68°C for 30s (extension), followed by 1 cycle at 68 °C for 480s (completion).

### FISH followed by immunofluorescence detection

All steps were conducted at room temperature unless otherwise indicated. 14 µm thick coronal cryosections were post-fixed for 5 min in 4 % PFA/PBS, 3 × 5 min washed in PBS (0.1 %, diethyl pyrocarbonate [DEPC]) and 1 × 5 min in 1x saline-sodium citrate (SSC, 0.1 % DEPC). Pre-hybridization was performed with 200 µL hybridization buffer (50 % formamide, 5x Denhardt’s solution, 5x SSC, 0.25 mg/mL yeast tRNA, 0.2 mg/mL salmon sperm DNA, DEPC-H2O) per slide for 4 hrs. Hybridization was performed with 1 ng/µL digoxigenin (DIG)-labelled probes overnight at 65 °C. Probes corresponded to 1) KCHIP4a: NT 1-192 of Mus musculus KCNIP4, transcript variant 4 (NCBI NM_030265.3); 2) KCHIP4 all: NT 455-839 of Mus musculus KCNIP4, transcript variant 4 (NM_001199242.1). Digoxygenin-labeled antisense probes were generated following standard procedures. Post-hybridization washes were performed at 60°C for 5 min in 5x SSC, followed by 5 min in 2x SSC and 20 min in 0.2x SSC/50 % formamide. Slides were allowed to cool down for 20 min and washed twice in 0.2x SSC. Endogenous peroxidase activity was quenched by incubation in 3 % H_2_O_2_/1x SSC for 15 min. Slides were washed twice for 5 min in TBS 1x followed by 30 min blocking in TNB blocking buffer (0.1 M Tris-HCl, 0.15 M NaCl, 0.5 % (w/v) TSA blocking reagent (PerkinElmer)). Slides were incubated in anti-DIG-POD Fab fragments (1:500; Roche) diluted in TNB buffer for 2 hrs. in a humidified chamber followed by 3 × 10 washing in TBS-T (0.1 % Tween-20). Tyramide signal amplification was performed with 100 µL TSA® Plus FITC (1:60; PerkinElmer) per slide for 10 min in a dark chamber. Sections were incubated with Ar6 buffer (1:10; PerkinElmer, Massachusetts) for 45 min in a heat steamer and allowed to cool down for 30 min.

Immunohistochemistry was performed following standard procedures. Briefly, slides were blocked for 30 min in 0.3 % Triton / 10 % normal goat serum / TBS followed by overnight primary antibody incubation (1:1000; rb TH; Merck Millipore) at 4 °C. Slides were washed 3 × 10 min in TBS-T and secondary antibody incubation (1:750; Life Technologies) was performed for 1 hr. After 3 × 10 min washing in TBS-T, slides were incubated in DAPI 1x for 10 min, washed 3 × 5 min with TBS and embedded in ProLongTM Gold Antifade Mountant (Thermo Fisher Scientific). Images were taken with a Nikon Eclipse90i microscope and acquisition was performed with the NIS Elements software (Version 5.01). Brightness and contrast were moderately enhanced in Photoshop across the entire image, and no further image processing was performed.

### Kv4 channel complex high-resolution proteomics

#### Crude membrane preparation and protein solubilization

Isolated cerebellum and midbrain samples of six CTRL and six Ex3d mice were individually processed. Tissues were homogenized two times in 0.5 mL homogenization buffer (320 mM Sucrose, 10 mM Tris/HCl, 1,5 mM MgCl2, 1 mM EGTA, 1 mM Iodoacetamide, pH 7.5) supplemented with protease inhibitors (Aprotinin, Pepstatin, Leupeptin (each at 2 µg/mL) and 1 mM PMSF) with a Dounce homogenizer. Homogenates were centrifuged at 100xg for 5 minutes, resulting supernatants centrifuged at 200.000xg (S45A rotor, Sorvall) for 20 minutes. Pellets were re-suspended in 0.5 mL Lysis-buffer (5 mM Tris/HCl, 1 mM EDTA, 1 mM Iodoacetamide, protease inhibitors) and after 30 minutes incubation time on ice centrifuged at 200.000xg for 20 minutes. The resulting pellets were re-suspended in 0.1 mL 20 mM Tris/HCl pH 7,5 and protein concentrations determined by Bradford assay. 0.3 mg of each sample were solubilized in 0.5 mL CL-48 buffer (Logopharm GmbH, Germany) with freshly added protease inhibitors for 30 minutes on ice. Non-solubilized material was removed by ultracentrifugation (8 min. at 125.000xg, S45A).

#### Affinity-purifications

Solubilisates were directly incubated with 7.5 µg Kv4.3 antibodies (Alomone, #APC-017) immobilized on Protein A Dynabeads (Invitrogen) for 2 hours at 4°C. Two additional 0.5 mL solubilisates of cerebellum and midbrain samples were incubated with 7.5 µg IgG (Millipore, 12-370), which were used as internal specificity control. After two washing steps with 0.5 mL CL-48 buffer bound proteins were eluted with 1x Laemmli buffer without DTT. Proteins were shortly run on 10 % SDS-PAGE and silver-stained. Samples were further processed as described ^20^, lanes were split in two pieces and digested with sequencing-grade modified trypsin (Promega, #V5111) and alpha-lytic protease (Sigma-Aldrich, #A6362). Peptides were extracted and prepared for MS analysis.

#### Mass spectrometry

Mass spectrometric analyses of peptide mixtures were carried out on a Q Exactive HF-X mass spectrometer coupled to an UltiMate 3000 RSLCnano HPLC system (both Thermo Scientific, Germany) as described ^69^. For each LC-MS/MS dataset, a peak list was extracted from fragment ion spectra using the “msconvert.exe” tool (part of ProteoWizard; http://proteowizard.sourceforge.net/; v3.0.6906; Mascot generic format with filter options “peakPicking true 1-” and “threshold count 500 most-intense”) and the precursor m/z values were shifted by the median m/z offset of all peptides assigned to proteins in a preliminary database search with 50 ppm peptide mass tolerance. Corrected peak lists were searched with Mascot (Matrix Science, UK) against all mouse, rat, and human entries of the UniProtKB/Swiss-Prot database (supplemented with mouse KChIP4 splice isoforms, identifier: Q6PHZ8-2, Q6PHZ8-3, Q6PHZ8-4, Q6PHZ8-5, Q6PHZ8-6). Acetyl (Protein N-term), Carbamidomethyl (C), Gln->pyro-Glu (N-term Q), Glu->pyro-Glu (N-term E), Oxidation (M), and Propionamide (C) were chosen as variable modifications, peptide and fragment mass tolerance were set to ± 5 ppm and ± 0.8 Da, respectively. One missed cleavage was allowed. The expect value cut-off for peptide assignment was set to 0.5.

#### Protein quantification

Label-free quantification of proteins was done as described previously ^21,22^. Peptide signal intensities (peak volumes, PVs) were determined and offline mass calibrated using MaxQuant (http://www.maxquant.org) and then assigned to peptides based on their m/z and elution time obtained either directly from MS/MS-based identification or indirectly (i.e., from identifications in parallel datasets) using in-house developed software. Molecular abundances of proteins were estimated using the abundance_norm, spec_ score defined as the sum of all assigned and protein isoform-specific PVs divided by the number of MS-accessible protein isoform-specific amino acids (Bildl et al., 2012). In order to visualize molecular compositions of Kv4-KChIP complexes normalized molecular abundance values of Kv4.2, Kv4.3 and KCHIP1, 2, 3 and 4, and DDP6 and DPP10, were calculated (Figure 2B and C, and Figure S2). Normalized values were obtained by dividing abundance_norm, spec_ values of individual proteins by the factor (∑ of Kv4 abundances/4). Hence, the sum of normalized molecular abundance values for Kv4.2 and Kv4.3 is 4. Relative abundances of KChIP4a isoform in Ex3d versus CTRL samples were determined by use of PV values of 3-5 isoform-unique peptide features. For each unique peptide, the PVs were first normalized to their maximum over 12 AP datasets from cerebellum and over 12 AP datasets from midbrain, yielding relative peptide profiles. Accordingly, relative abundance differences of KChIP4a were determined as mean value of peptide profiles from six Ex3d samples and six CTRL samples from cerebellum and midbrain respectively, whereby to the CTRL was set the value of 1 (Figure 2D).

### Patch-clamp recordings of projection-identified DA neurons

#### Retrograde tracing

Mice were premedicated with Carprofen (Rimadyl, Pfizer, Berlin, Germany, 5 mg/kg s.c.). Anesthesia was induced and maintained throughout the surgery using isoflurane (AbbVie, North Chicago, USA) Induction: 4 % in 350 mL/min O2, Maintenance dose: 1-2.5 % in 350 mL/min O2). For local anesthesia and analgesia, we administered a mixture of Lidocaine and Pirlocaine (EMLA, Aspen, Munich, Germany). After induction of anesthesia, mice were transferred in a stereotaxic frame (Kopf Instruments, Tujunga, CA, USA) where 100nl of red retrobeads (diluted 1:30 in aCSF, Lumafluor, Naples, Florida, USA) were injected in different areas of the striatum (in reference to Bregma: NAcc core: AP: 1.54 mm, ML: ±1.0 mm, DV: 4.0 mm; NAcc lateral shell: AP: 0.86 mm, ML: ±1.75 mm, DV: 4.5 mm; DLS: AP: 0.74 mm, ML:± 2.2 mm, DV: 2.6 mm) to label midbrain DA neurons, as previously described ^4^.

#### *Ex vivo* electrophysiology

After three days of recovery, to ensure sufficient retrograde axonal transport of the red retrobeads, the mice were perfused transcardially using ice cold perfusion solution (125 mM NaCl, 2.5 mM KCl, 6 mM MgCl_2_, 0.1 mM CaCl_2_, 25 mM NaHCO_3_, 1.25 mM NaH_2_PO_4_, 50 mM Sucrose, 2.5 mM Glucose, 3 mM Kynurenic acid, bubbled with Carbogen). Brains were quickly removed, and the midbrain was sliced into 250 µm thick coronal slices using a vibratome (Leica VT1200S, Leica Biosystems, Wetzlar, Germany). Slices were then transferred to a beaker containing ACSF (125 mM NaCl, 3.5 mM KCl, 1.2 mM MgCl_2_, 1.2 mM CaCl_2_, 25 mM NaHCO_3_, 1.25 mM NaH_2_PO_4_ constantly bubbled with carbogen) and allowed to recover at 37°C for 60 minutes.

After recovery, slices were transferred to a recording chamber and constantly perfused at a rate of 2-4 mL/minute^-1^ with ASCF maintained at 37 °C (Temperature controller VI, Luigs Neumann, Ratingen, Germany). Synaptic transmission was blocked using CNQX (12.5 μM, Biotrend), DL-AP5 (10 μM, Tocris) and Gabazine (4 μM, SR95531, Biotrend). Midbrain DA neurons were visualized using a light microscope (Axioskope 2 FS plus, Zeiss, Oberkochen, Germany) and an infrared camera (VX55, TILL photonics). Red retrobeads were excited by an epifluorescence lamp (HBO100 Nikon) filtered at 546/12 nm. DA neurons containing red retrobeads were recorded using borosilicate pipettes (3-4 MOhm, GC150TF, Harvard Apparatus) filled with internal solution (135 mM K-gluconate, 5 mM KCl, 10 mM HEPES, 0.1 mM EGTA, 5 mM MgCl_2_, 0.075 mM CaCl_2,_ 5 mM ATP, 1 mM GTP, 0,1 % Neurobiotin, pH 7.35 (290-300 mOsmol). Data were recorded at 20 kHz and filtered with a low-pass filter Bessel characteristic of a 5 kHz cut-off frequency using an EPC 10 amplifier (HEKA Elektronik, Lambrecht, Germany). Patchmaster (HEKA Elektronik, Lambrecht, Germany), IgorPro (WaveMetrics, Portland, Oregon, USA) and MATLAB (MathWorks, Massachusetts, USA) were used for acquisition and analysis. Only spontaneously active cells that showed robust pacemaking, and that were post-hoc identified as TH positive and containing red retrobeads, were included in this study.

#### Histology

The forebrain tissue-block containing striatal injection sites, as well as midbrain slices containing the recorded and neurobiotin-filled neurons, were immersion-fixed at 4°C overnight in a solution of 4 % paraformaldehyde and 15 % picric acid in phosphate buffer solution (PBS). Striatal injection sites were cut into 100 µm coronal sections using a vibratome (VT1000S, Leica Biosystems, Wetzlar, Germany). Free-floating sections were washed in PBS and incubated with blocking solution (10 % horse serum, 0.5 % Triton X-100 % and 0.2 % BSA in PBS) at room temperature for 1 h. Afterwards, sections were incubated in carrier solution (1 % horse serum, 0.5 % Triton X-100 % and 0.2 % BSA in PBS) containing the primary antibody (polyclonal rabbit anti-TH, 1:1000, Synaptic Systems Cat# 213 104) overnight at room temperature. On the second day, sections were washed several times in PBS and incubated with carrier solution containing the secondary antibody (goat anti-rabbit 488, 1:750, Thermo Fisher Scientific Cat# A-11011) and in case of the midbrain sections Streptavidin AlexaFluor 408 (1:750, Invitrogen) overnight at room temperature. On the third day, sections were washed in PBS, mounted, and stored at 4°C.

### Biophysical modelling of DA neurons

#### Model calibration

The A single compartment, core projecting DA neuron model was adapted from Knowlton et al. (2021). The inactivation variable of the Kv4.3 channel model from that study was separated into an additive combination of slow and fast variables. Steady state activation, inactivation, and kinetics of inactivation were set to be consistent with voltage clamp data in Figure 4 and are given in Supplemental Table 1. Activation kinetics were not altered from those originally adapted from Tarfa et al (2017) and was assumed to be identical for channels with slow or fast inactivation gates. Steady state activation and inactivation are given by Boltzman functions while the inactivation time constants are given by Boltzman sigmoids between indicated minimum and maximum values. The relative contributions of the slow and fast Kv4.3 channels, determined by the binding of different auxiliary subunits, is the only intrinsic model parameter varied between CTRL and Ex3D models.

The model was implemented in NEURON ^70,71^ with extensions for parallel computing ^72^. Model files are freely available at https://senselab.med.yale.edu/modeldb/ShowModel?model=267500 (password: okaychip).

#### Balanced state methods

A simulated *in vivo* balanced state was implemented by simultaneous application of high frequency, low conductance (100 Hz) Poisson GABA_A_ and NMDA receptor activation ^29,73–75^. The NEURON implementation of NMDA receptor activation was taken unmodified from Moradi et al. (2013). GABA_A_ receptor activation uses NEURON’s built in bi-exponential synapse with a rise time of 0.2 ms and a fall time of 2.4 ms. The reversal potential of chloride was fixed at -60 mV for all balanced state GABA_A_ and the reversal potential of NMDA at 0 mV. The Ca^2+^ influx via NMDA channel was not included in the pool that activates the SK K+ current. Balanced state synaptic weights for NMDA and GABA_A_ were set to be constant. These weights were chosen to obtain a simulated *in vivo* firing rate was consistent with *in vivo* recordings, and the GABA_A_ conductance was of sufficient magnitude to silence cells in the absence of excitation, consistent with a balanced state.

Inhibitory synaptic inputs, as would putatively underlie the signaling of negative RPES, were modelled as additional GABA_A_ inputs applied synchronously. Inhibition trials were repeated then recombined into a histogram of simulated network response under an ergodic approximation. Each histogram was constructed from 500 simulated trials. An inter-trial refractory period of 4s was used to eliminate transients. Multi-threading was used to run different parameterizations of the RPE trial.

### Behavior

#### Open field

Spontaneous locomotor and exploratory activity (track length, wall distance, time in center and number of rearings) were evaluated in an open field arena (a lidless box measuring 52 × 52 cm, under red illumination at 3 lux) using a video tracking system (Viewer II, Biobserve, Germany) as described in a previous study from our group ^77^. For a comprehensive phenotyping of mutant mice, the total track length, the time spent and track length within the center area of the arena (defined as a 30×30 cm square zone with all sides equidistant from the walls), as well as the number and duration of rearings, were evaluated. Rearings were recorded via infra-red beam breaks at a height of 4.5 cm and defined by being at least 200 ms long and two subsequent rearings had to be at least 80 ms apart. After ten minutes, the mice were returned to their home cages for at least two minutes, and then subjected to the novel object recognition test.

#### Novel object recognition

A novel object exploration and preference task was used to assess the mice’s recognition, memory retention and preference for novel stimuli ^78,79^. In this task, mice were placed in the same open field arena described in the previous section, after the open field test (which also served as a habituation for the novel object recognition test), but within the arena two identical objects (stainless steel cylinders; 3 cm diameter × 6 cm height) were placed at equal lengths from each other and at 15 cm from the upper left and right corners. The mice were allowed to freely explore the arena and the objects for 10 minutes (trial 1), after which they were removed from the arena. Subsequently, one of the objects was replaced by a different, novel object (plastic coated rectangular prism; 3×3 cm base × 6 cm height), and the mice were again allowed to explore the arena and the objects (trial 2).

Object recognition was analyzed using Biobserve’s Object Recognition plug-in, and object interaction events were defined as periods in which a mouse was directly facing the object (snout directed to the object within a 180° angle) at a distance shorter than 3 cm. Both the number and duration of these interaction events were quantified for both trials of the task. Exploration dynamics (number and duration of objected-directed exploration) were analyzed for both trials, as well as the differences between exploration of an already explored (“old”) and novel (“new”) object in trial 2, were analyzed. For the quantification of novel object discrimination and preference, a discrimination index was calculated by dividing the difference between the time spent exploring the novel object and the time spent exploring the familiar object by the sum of these two measures ^78,79^.

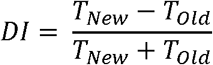

where

*DI* = discrimination index

T*_New_*= time spent exploring the novel object

*T_old_*= time spent exploring the familiar object

#### Hole board

Exploration of holes, a naturalistic behavior for mice, was evaluated with the hole board task ^80^. In this task, mice were placed for five minutes in an open field arena similar to the one used for the open field test, but with 4 circular holes (2 cm in diameter) on its floor (Actimot2, TSE systems). When confronted with such an arena, mice tend to spontaneously check the holes by dipping their heads in them. The performance of head dips was recorded with infra-red beam breaks placed immediately under the inferior surface of the floorboard. The latency to the first head dip, as well as the total number and duration of head dips, were quantified. The percentage of repeated head dips (two dips performed sequentially into the same hole) as well as the percentage of dips into the preferred hole (the hole that was most explored during the session) were also quantified and compared between genotypes.

#### Spontaneous alternation in the plus maze

Working memory performance was quantified between genotypes using the spontaneous alternation in the plus maze ^81^. Mice were placed in a plastic plus maze (four equidistant 35 × 4.5 cm arms radiating at 90° angles from a circular central arena with 10 cm diameter; walls were 15 cm in height) and allowed free exploration for 12 minutes ^82^. An arm entry was defined as when the mouse enters an arm with all its four paws. Each session was recorded with a video camera and quantification of arm entries was made through visual analysis of the videos. A spontaneous alternation was marked when the mouse explored four different arms in five consecutive arm entries, and the proportion of spontaneous alternations was quantified as the number of real alternations divided by the total possible number of alternations (sum of all arm entries minus four); chance performance in this task is calculated to be 44 % ^81,82^.

#### Reinforcement learning task

Male and female Ex3d (N = 10) and CTRL (N = 20) mice, over 8 weeks old, were genotyped and individually identified by the implantation of a subcutaneous transponder microchip (1.4 mm × 9 mm ISO FDX-B glass transponder, Planet ID, Germany). Behavioral comparisons were performed between Ex3d and DAT-Cre KI mice. One Ex3d mouse showed erratic behavior within the conditioning box and was excluded from data analysis (final N of Ex3d group = 9). Mice were motivated by water restriction (∼85 % of their initial body weight) and were rewarded with a solution of 10 % sucrose in tap water. Daily water rations varied between 1 and 1.5 mL depending on the mice’s weight on that day, with a set target of 85 % of the initial body weight. Except during experimental sessions, water was always delivered in a cup placed in their home cage. Mice were also closely monitored in order to ensure that the water supply was consumed and not spilt over or contaminated and the health of the water restricted mice was also evaluated daily ^83^. This protocol maintained a stable body weight around the desired target during all experimental sessions (Figure S4) and did not result in signs of overt dehydration or ill health (e.g., hunched posture or ruffled fur).

The full experimental paradigm spanned 24 days. On the first four days, mice were submitted only to water restriction. In the following two days, they were tamed, i.e., gently handled (held up above their home cage on a spread palm, with no constriction or entrapment by the experimenter) until they no longer tried to escape from the experimenter’s hand, showed no overt signs of stress and anxiety and readily drank a portion of liquid reward (0.2 mL) given by the experimenter via a syringe while being held. The following day, the mice were placed inside the operant chamber with the reward port removed for 50 minutes in order to acclimate to the experimental conditions. After this time, they were returned to their home cage and given their daily ration of water. The day after that, mice underwent shaping, i.e., they were placed in the operant chamber for another 50 minutes, now with the reward port present, and at semi-random time intervals (mean of 60 seconds, varying between 30 and 90 seconds), a reward portion (16.66 µL) was delivered at the port, with no cue to its delivery. The following day, mice were submitted to the conditioning task.

The task used in this study is based on the study by Steinberg et al. (2013). In this paradigm, mice learned to associate a CS (sound tone pulsed at 3 Hz – 0.1 seconds on/0.2 seconds off – at 70 dB) to the availability of reward in the reward port. Each session consisted of ten trials (mean inter-trial interval of 4 minutes, varying between 1.5 minutes and 6.5 minutes) in which the auditory cue was on for 30 seconds. Mice could trigger reward delivery to the port by entering it during CS presentation. Rewards were delivered in a cycle of 2 second reward delivery (16.66 µL) followed by a 3 second consumption interval. Delivery was continuous for as long as the mouse kept its head in the port during CS presentation. This allowed for a maximum of 6 rewards per trial and a maximum of 60 rewards per session (a total of ∼1 mL or reward in the task per day). Additional water supplementation, when needed to complete the mice’s daily water ration, was provided in the home cage as described in the previous paragraph.

This acquisition phase lasted for 11 daily sessions. After acquisition, extinction of the conditioned response was tested by consistently omitting the reward during CS presentation for six daily sessions. After extinction, mice were returned to their home cage and received water ad libitum for at least three days before being submitted to other behavioral tasks. All mice recovered their initial body weight in this period of time and showed no long-term adverse effects from water restriction.

Performance in this task was quantified by the total time the mice spent in the port during the cued trials and the latency to enter the port after cue onset. Time in port was normalized both as a percentage of total reward availability time and also by subtracting the amount of time the mouse spend in the port 30 second before cue onset ^17^. This means that CS-US associations were quantified by the mouse responding both quickly and selectively to the cue. In addition, the time stamps of every head entry and head retraction into the port were recorded, which allowed the quantification of the dynamics of head entries during cue presentation and during the ITI. Latency was quantified as the time between cue initiation and the first head entry into the reward port; if mice did not respond in a trial, the maximal possible latency value (30 s) was ascribed to that trial. The dynamics of head entries during extinction were used to infer how the mouse adapts its behavioral response in the absence of an expected reinforcement, i.e., whether it perform fewer head entries (a proxy of the initiation rate of reward-seeking behaviors), head entries that are just shorter in length (indicating a faster disengagement from reward-seeking behaviors) or a combination of both.

### Behavioral modelling

A modified Rescorla-Wagner model, which applied two different learning rates for positive (α_P_) and negative (α_N_) RPEs, was used in order to formally quantify differences in learning between Ex3d and CTRL genotype groups ^31–33^. In detail:

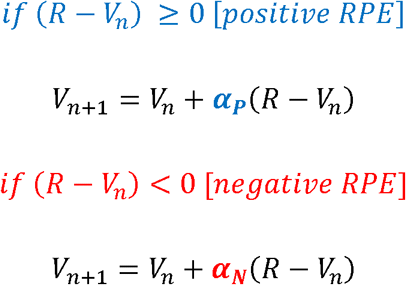

where

*V* = associative strength between CS and US

α_*P*_ and α_*N*_ = learning rates from positive and negative RPEs, respectively

*R* = maximal associative strength of the US (reward value)

*n* = trial index

As a control, the same model was also applied assuming a single learning rate for both positive and negative RPEs, i.e., a classical Rescorla-Wagner model (Figure S5). All modelling and fitting were done with custom MATLAB® code. The percentage of time in port metric was assumed to be a linear read-out of *V*. The limits of *V* in relation to the empirical data were bound by the initial value of *V* before the task (*V_0_*), the lower and upper limits of associative strength readout (*V_min_* and *V_max_*, respectively). These parameters were derived empirically from each mouse, with *V_0_* being set as the average response in the first three trials of acquisition, and *V_min_* and *V_max_* being set as the average responses in the last two sessions of acquisition and extinction, respectively. Fitting of learning rates was performed with least absolute residuals method by implementing the *fminsearch* MATLAB function ^31^. Goodness of fit was inferred from the residual values of the optimal fit for each mouse.

### Statistical analyses

For all two-group comparisons, data were tested for normality with the Kolmogorov-Smirnov test. If both distributions were Gaussian, differences were analyzed using two-tailed T-tests; otherwise, the two-tailed Mann-Whitney test was used. Comparisons between more than two groups were tested using one-way ANOVA. Comparisons between groups over multiple trials or sessions were evaluated using two-way repeated measures ANOVA with Sisak’s multiple comparisons *post-hoc* tests. Statistical significance was set at *P* < 0.05 for all comparisons.

Figures were created with MATLAB, GraphPad Prism 8, and BioRender.com.

## Supplemental tables

**Table S1.**
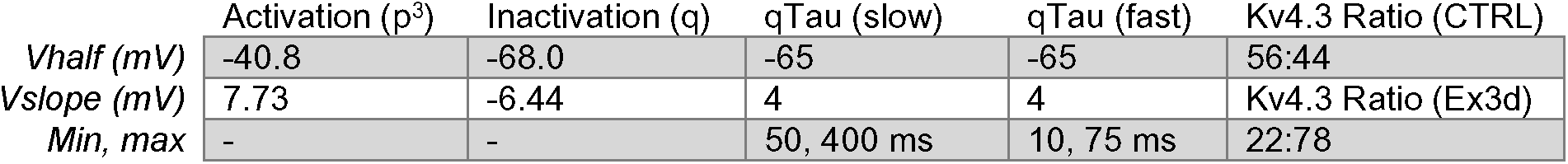
Steady state activation, inactivation, and kinetics of inactivation of the Kv4 currents of the biophysical models of atypical DA neurons, related to Figure 5.

| | Activation ($p^3$) | Inactivation ($q$) | qTau (slow) | qTau (fast) | Kv4.3 Ratio (CTRL) |
| --- | --- | --- | --- | --- | --- |
| *Vhalf (mV)* | -40.8 | -68.0 | -65 | -65 | 56:44 |
| *Vslope (mV)* | 7.73 | -6.44 | 4 | 4 | Kv4.3 Ratio (Ex3d) |
| *Min, max* | - | - | 50, 400 ms | 10, 75 ms | 22:78 |

## Supplemental figures

**Figure S1.**
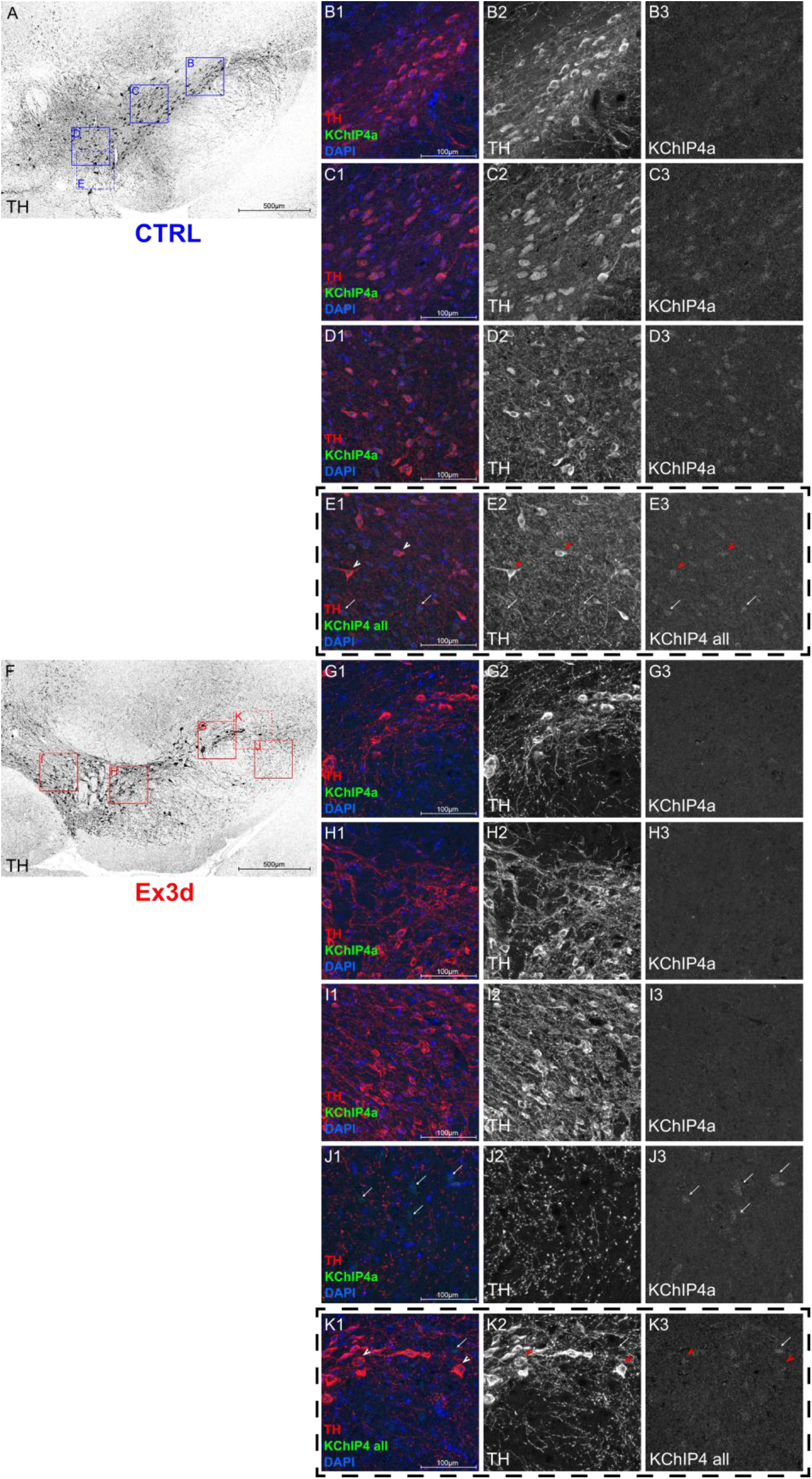
FISH assay confirms that Ex3d mice have a selective ablation of only KChIP4a mRNA expression specifically in DA neurons, related to Figure 1. **A:** Overview of a midbrain slice from a CTRL mouse where the indicated rectangles represent the ROIs used for panels B-E. **B-C:** FISH labeling with the KChIP4a-specific probe reveals that CTRL mice show robust expression of KChIP4a mRNA in TH^+^ and TH^-^ neurons across the whole midbrain. **E:** Labeling with the general KChIP4 probe, which binds to all KChIP4 isoforms, also shows expression in midbrain TH^+^ and TH^-^ neurons. **F:** Overview of a midbrain slice from an Ex3d mouse where the indicated rectangles represent the ROIs used for panels G-K. **G-I:** unlike in the CTRLs, labeling with the KChIP4a-specific probe showed that there was no expression of KChIP4a in TH^+^ neurons across the whole midbrain of Ex3d mice, including the SN and VTA. **J:** However, we did observe robust KChIP4a expression in midbrain TH^-^ neurons, demonstrating that the removal of KChIP4a was cell-type specific. K: Labeling with the general KChIP4 probe showed expression of KChIP4 variants in TH+ neurons in the midbrain of Ex3d mice, demonstrating that the removal of KChIP4a was also splice variant-specific.

**Figure S2.**
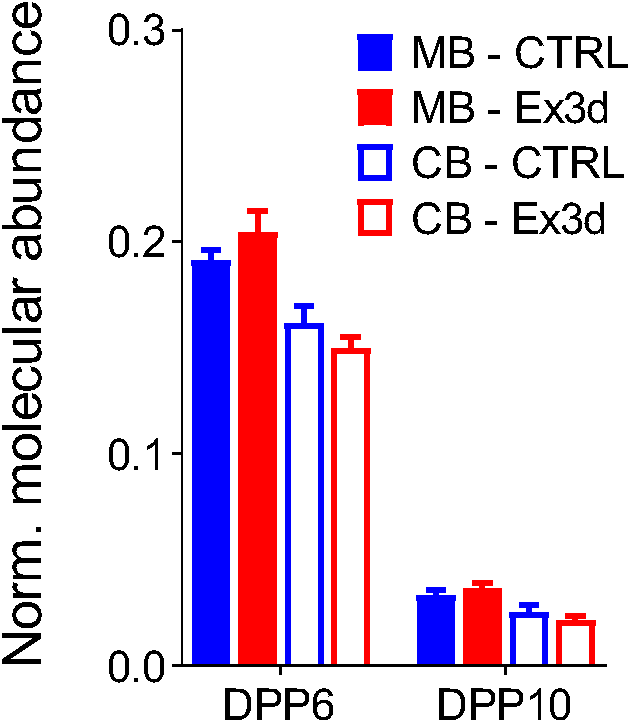
Relative abundance of DPPX proteins bound to Kv4 channels in the midbrain and in the cerebellum, related to Figure 2. We found relatively low abundance of DPP6 and DPP10 proteins bound to Kv4 channel complexes in both the MB and CB. Importantly, there was no effect of the Ex3d mutation in either region. Error bars indicate SEM.

**Figure S3.**
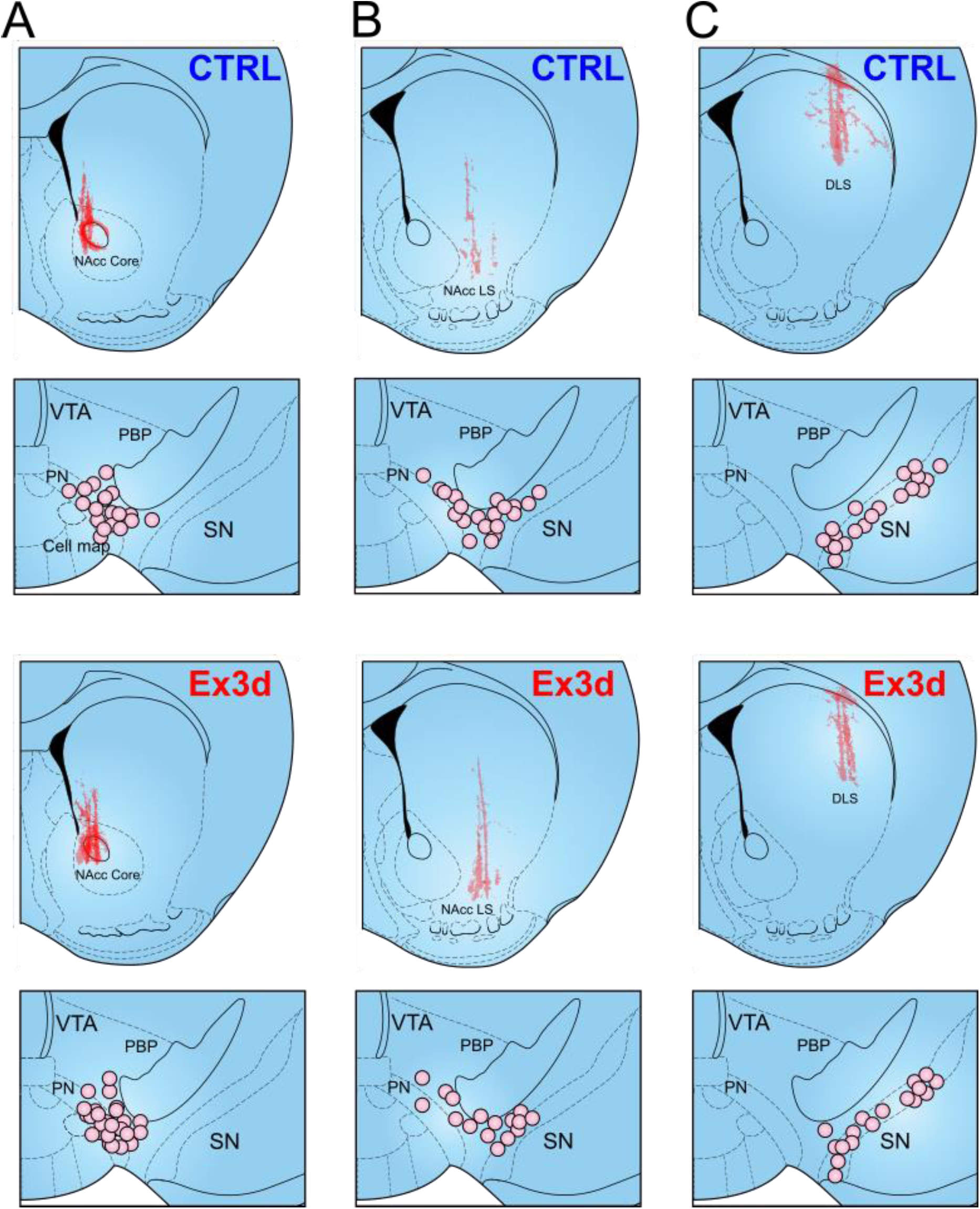
red retrobead injection sites and recording locations for projection-specific patch-clamp experiments, related to Figure 3. **A,B,C:** Spread of all red retrobead injections and location of projection-identified TH+ cells in the midbrain for the DA-cNAcc (A), DA-lNAcc (B) and DA-DLS (C) neuronal populations in CTRL and Ex3d mice.

**Figure S4.**
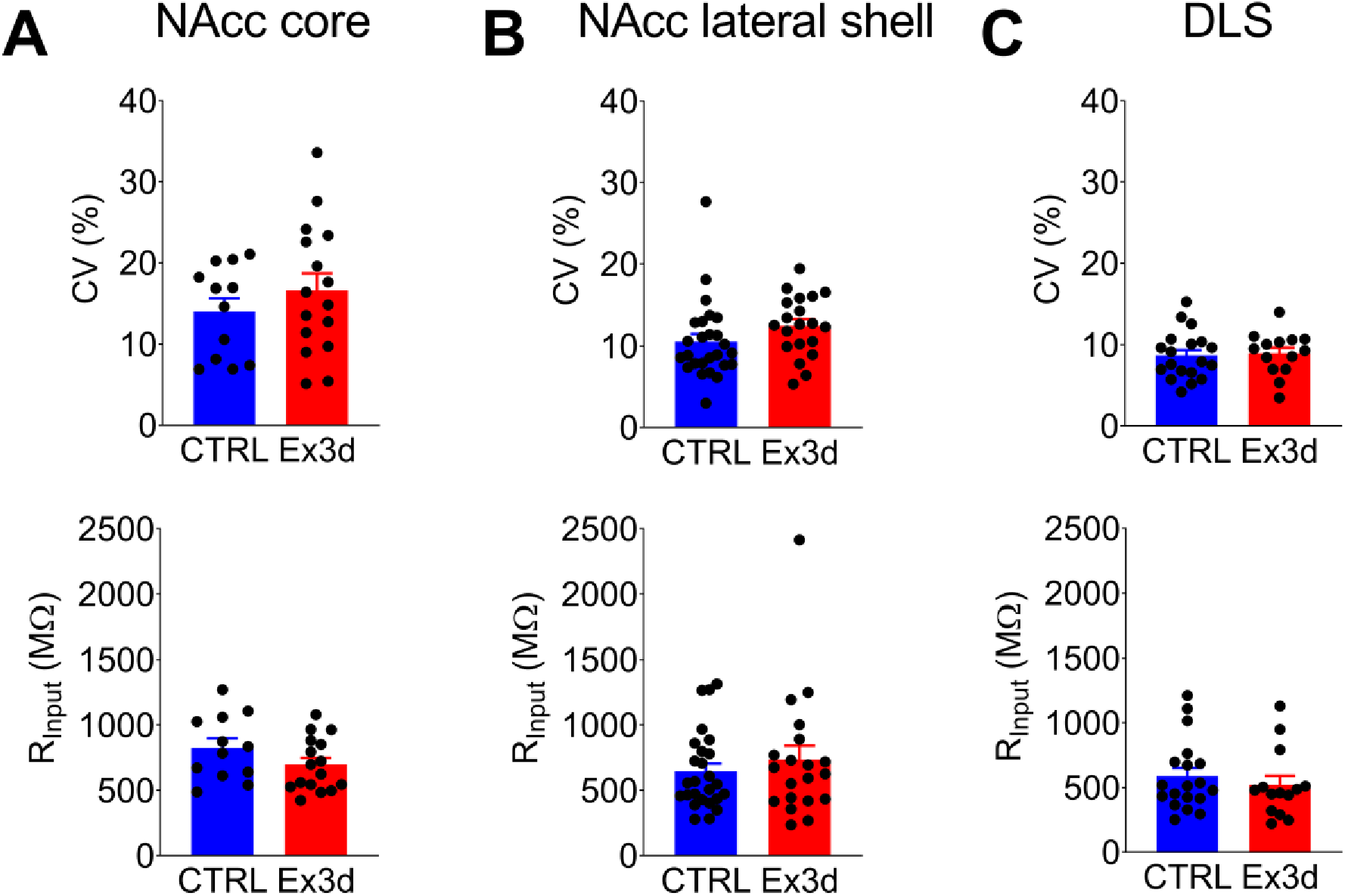
Additional biophysical properties of the recorded DA neuron subpopulations are not affected by the Ex3d mutation, related to Figure 3. **A:** Values of input resistance (R_input_) and inter-spike-interval coefficient of variation (CV) for DA-cNAcc projecting neurons of CTRL and Ex3d mice. **B,C:** same as above, but for DA-lNAcc and DA-DLS neurons. Note there were no significant genotype effects for any of these variables in any of the projection-identified populations. Error bars indicate SEM.

**Figure S5.**
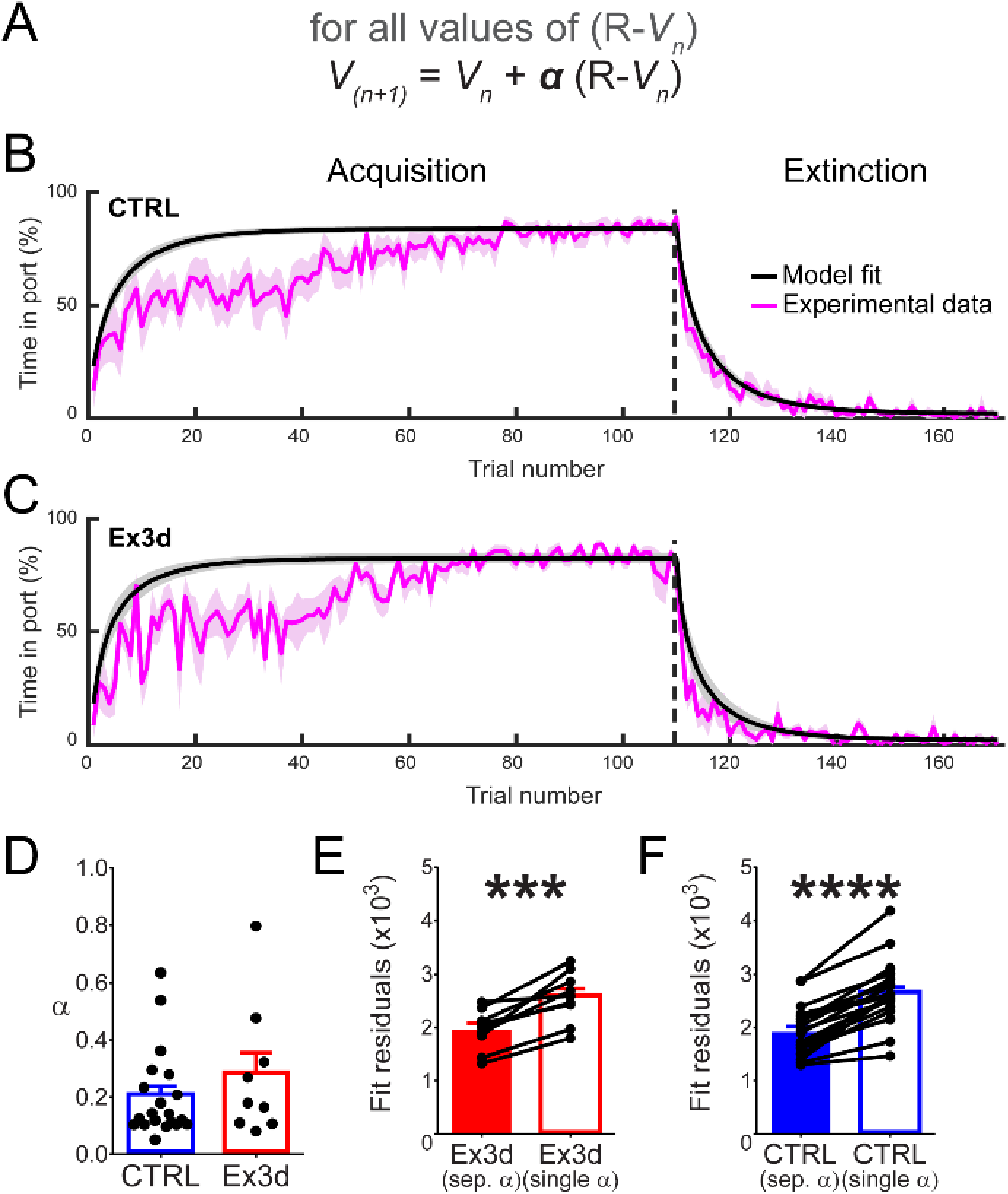
Models that do not assume different learning rates for positive and negative RPEs produce significantly worse fits and do not replicate the observed Ex3d behavioral phenotype, related to Figure 6. **A:** Equation of the traditional Rescorla-Wagner model used as a control. **B,C:** Comparison of model fits and experimental data for the CTRL (B) and Ex3d (C) group. **D:** fitted learning rates for CTRL and Ex3d data. Note the lack of genotype differences. **E, F:** comparison of fit residuals between models with a single learning rate and with separate positive and negative learning rates for the Ex3d (E) and CTRL (F) groups. Note that single learning rate models are significantly worse at fitting the behavioral data. Error bars indicate SEM.

